# Modeling spatial, developmental, physiological, and topological constraints on human brain connectivity

**DOI:** 10.1101/2021.09.29.462379

**Authors:** S. Oldham, B. D. Fulcher, K. Aquino, A. Arnatkevičiūtė, C. Paquola, R. Shishegar, A. Fornito

## Abstract

The complex connectivity of nervous systems is thought to have been shaped by competitive selection pressures to minimize wiring costs and support adaptive function. Accordingly, recent modeling work indicates that stochastic processes, shaped by putative trade-offs between the cost and value of each connection, can successfully reproduce many topological properties of macroscale human connectomes measured with diffusion magnetic resonance imaging. Here, we derive a new formalism with the aim to more accurately capture the competing pressures of wiring cost minimization and topological complexity. We further show that model performance can be improved by accounting for developmental changes in brain geometry and associated wiring costs, and by using inter-regional transcriptional or microstructural similarity rather than topological wiring-rules. However, all models struggled to capture topologies spatial embedding. Our findings highlight an important role for genetics in shaping macroscale brain connectivity and indicate that stochastic models offer an incomplete account of connectome organization.

## 1 Introduction

The human brain is a topologically complex network, showing properties that are neither completely random nor completely regular (*1*). These properties are commonly studied through the lens of graph theory (*1, 2*), in which the brain is represented as a collection of nodes, depicting putative processing units, such as individual neurons, neuronal populations, or brain regions, linked by edges, that capture some aspect of structural or functional connectivity between nodes. The application of graph-theoretic tools has identified key topological properties of brain networks, such as the existence of highly connected network hubs, a rich-club of strong inter-connectivity between hubs, and an economical, small-world, hierarchically modular organization (*1–4*). These topological properties also a have a characteristic topography, meaning that they are spatially embedded in consistent locations; for instance, network hubs tend to be located in transmodal association cortices (*4, 5*). Understanding the causes and consequences of this complex arrangement of connections is a central objective of connectomics (*6*).

Over a century ago, Santiago Ramón y Cajal proposed some general principles for brain organization, arguing that nervous systems are configured according to three simple laws related to the conservation of space, material, and time (*7*). The conservation of space and material refers to a pressure to minimize the physical, metabolic, and cellular resources required to sustain neural function. This cost-minimization principle is ubiquitous in biological systems and minimizes unnecessary energy expenditure, which is essential for metabolically expensive organs such as the brain (*8*). Conservation of time refers to a requirement for rapid and efficient communication between system elements, which is essential for adaptive function and organism survival.

Several studies have suggested that cost-minimization is an important principle of neural organization, showing that properties as diverse as the spatial arrangement of neurons and cortical areas (*9–11*), the branching patterns of neuronal arbors (*12*), and the fraction of cortical grey matter occupied by axons and dendrites (*13*), can be explained by a pressure to minimize the overall volume of axonal wiring, which is often used to represent the wiring cost of a nervous system. However, a network configured solely to minimize wiring costs resembles a lattice, in which each element only connects to its nearest spatial neighbors. Abundant evidence indicates that connectomes have more long-range projections than expected under a pure cost-minimization model (*14–16*). These long-range projections are thought to act as topological shortcuts, improving the speed, efficiency, robustness, and complexity of communication across the network (*14, 16, 17*). In the language of Cajal, they can be said to conserve time. However, the long distances spanned by these connections (*5, 18*) connections incurs a wiring cost, leading to a trade-off between Cajal’s conservations laws; more specifically, between the conservation of space and material (wiring cost) on the one hand and the conservation of time or, more generally, the promotion of complex, adaptive processes (functional value), on the other (*16*).

Insight into the possible role of cost-value trade-offs in sculpting connectome topology has come from generative network models, which specify wiring rules for growing brain-like networks *in silico*. Empirical evidence has indicated that the probability that two neural elements (such as individual neurons or brain regions) are connected decays roughly exponentially as a function of the distance between them, termed ‘the exponential distance rule’ (EDR) (*19, 20*). Modeling studies indicate that it is possible to grow synthetic networks that capture many key topological properties of empirical connectomes according to this rule, when it is implemented as a stochastic process in which the probability of forming a connection between any two network nodes declines exponentially as a function of their anatomical distance (*19–23*). Under this purely spatial model, long-range connections are more costly than short-range connections, but wiring costs are not absolutely minimized and are subject to stochastic fluctuations around a characteristic connection length scale. The networks that result from this model show many complex topological properties identified in empirical connectome data, including modularity, a fat-tailed degree distribution, brain-like motif spectra and distributions of connection distances, and the presence of a densely-connected core (*19, 20, 24–26*). This exponential distance rule has thus been invoked as a fundamental principle of neuronal connectivity (Ercsey-Ravasz et al., 2013; Horvát et al., 2016; Wang & Kennedy, 2016).

Recent modeling of human connectome data suggests that the EDR offers an incomplete account of connectome architecture. Specifically, this work indicates that EDR-based models are less accurate in reproducing several topological features of empirical connectomes when compared to models that combine a distance penalty with a preference to form topologically favorable connections, thus more closely capturing the cost-value trade-off implicit in Cajal’s laws (*5, 28*– *32*). In particular, these studies suggest that models combining a distance penalty with a homophilic attachment rule, in which connections are more likely to form between nodes that connect to other similar nodes (*28–30, 33*), offer better accounts of the empirical data. However, three key considerations, detailed in the following, should be addressed before the homophilic attachment model can be accepted as a parsimonious account of macroscale human connectome topology.

A first consideration is that the models considered to date have quantified wiring costs at a single, post-natal timepoint, which does not account for the dramatic changes in brain size and geometry that occur during early development, when connections are being formed (*34, 35*). Between 18 weeks gestational age and birth, the brain undergoes an approximately 20-fold expansion in volume (*35*), which is coupled with substantial increases in the complexity of cortical folding (*34*). Such geometric changes influence distances between regions and may change inter-regional wiring costs when compared to the adult brain (*36, 37*), and modeling work has found that capturing the physical growth of biological neural networks is important to predicting their subsequent topology (*38, 39*). The effect of these changes in brain size and shape on model performance has not been considered; indeed, it is possible that even a simple EDR may provide a sufficient model if long-range connections are established early when distances are smaller and wiring costs are lower.

A second consideration is that current cost-topology models rely on abstract topological rules for influencing connection probabilities, which can sometimes have an ambiguous physiological interpretation. For instance, homophilic attachment based on similarity in connectivity neighborhoods implies that two nodes have knowledge of each other’s neighbors when forming a connection. It is unclear how such a mechanism would be instantiated in brain development. Alternative, physiologically-grounded homophilic processes may offer a more interpretable model. For instance, the architectonic type principle, formulated following extensive observations of mammalian tract-tracing data, proposes that regions with similar cytoarchitecture and laminar organization are more likely to be connected to each other (*40, 41*). Similarly, there is growing evidence that similarity in regional transcriptional profiles may also be linked to inter-regional connectivity (*5, 23*). Whether such physiologically-grounded homophilic attachment rules offer a better account of empirical data than topological homophily has not been evaluated.

A final consideration is that the performance of existing models is commonly evaluated with respect to topological properties of the network, while ignoring spatial topography. This is an important oversight, as the same topological distribution can be spatially embedded in different ways, and these topographical variations can have functional consequences (*42, 43*). An adequate model should ideally capture both topological and topographical properties of the empirical data. Recent evidence indicates that existing-cost-topology models cannot capture the topography of certain properties, such as the network degree sequence and, by extension, location of connectome hubs, even when model parameters are optimized for this objective (*5, 31, 32*) (although see also (*30*)).

Informed by these considerations, we use generative network models to investigate spatial, developmental, physiological, and topological constraints on the human connectome. First, we develop a framework to account for developmental changes in brain size and shape when estimating model-based wiring costs, thus yielding a new class of developmentally-informed models that offer a more realistic appraisal of how wiring costs shape connectome topology. Second, after introducing a new formulation of the cost-value model that more accurately captures trade-off mechanisms and which yields more interpretable parameter estimates, we compare the performance of spatial and trade-off models to models that rely on either topological or physiologically constrained wiring rules. Finally, we evaluate model performance with respect to both topological and topographical properties of the human connectome, yielding a comprehensive characterization of model performance.

## 2 Results

We used diffusion imaging data from 100 unrelated participants in the Human Connectome Project (*44*) to construct structural brain networks with which to assess the performance of different generative network models (see Methods). A schematic overview of our model fitting and evaluation procedure is presented in Fig. 1. As wiring costs are a fundamental element of the models that we evaluate, our first aim was to incorporate into the generative models the pronounced changes in cortical shape and size that occur during the second half of gestation, when most connections are being formed. To this end, we used cortical surface reconstructions of fetal structural MRI templates. These templates were obtained from a public database, where 81 scans of fetuses acquired between 19-39 weeks gestational age (GA) were used to construct templates spanning 21-38 gestational weeks (Fig. 1A) (*45, 46*). We registered each surface to an adult template surface using the Multimodal Surface Matching (MSM) algorithm (Fig. 1B) (*47, 48*), allowing us to map nodes to consistent spatial locations across all time points and to measure how distances between nodes, as a proxy for putative wiring costs, change through development. We refer to these developmentally informed models as ‘growth’ models and the traditional models that only estimate wiring cost in the adult brain as ‘static’ models (Fig. 1C).

**Fig. 1.**
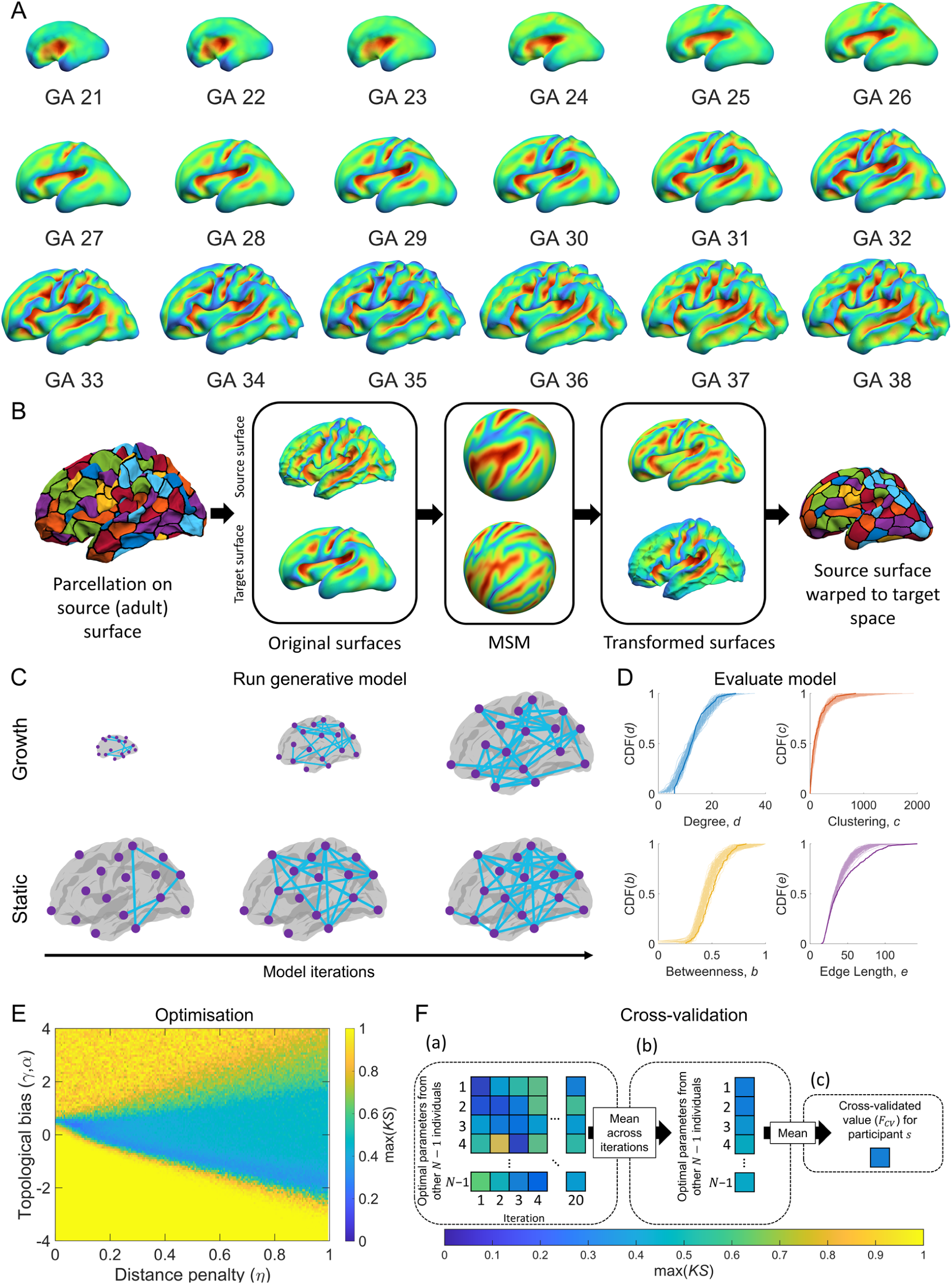
Schematic of our approach to fitting and evaluating generative connectome models. **(A)** Fetal surfaces from 21-38 weeks gestational age (GA), showing sulcal depth. **(B)** To estimate how network wiring costs change through development, we parcellate the adult surface and use spherical Multimodal Surface Matching (MSM) based on sulcal depth to map the parcellation to each of the fetal (target) surfaces. This procedure allows us to track the location of each parcellated region and estimate how wiring costs change through development. **(C)** Generative models were run using estimates of wiring cost based on adult distances (static models) or distances that change over time (growth models). **(D)** Benchmark topological properties were measured for each synthetic network and compared to the distributions of the empirical network using the *KS* statistic to quantify model fit (similar distributions indicate a better fit; thick lines represent empirical data and lighter lines correspond to different realizations of a model). **(E)** Steps **C**-**D** were repeated for different parameter values in each model using an optimization scheme that searches the parameter landscape to find the parameter combination that yields the best fit to the data. Starting with an initial random sample, the algorithm narrows in on areas of the landscape associated with better fits and samples those regions more often (see Methods). **(F)** Leave-one-out cross-validation is used to avoid over-fitting. For a given participants network, the best-fitting parameters for the 99 other participants are used to generate model networks. These models are iterated 20 times to account for the inherent stochasticity of the models (a) and the average across these 20 iterations is then taken to yield 99 fit values (b). These 99 fit values are the averaged (c), resulting in a single, cross-validated fit statistic, *F*_*CV*_ for each person.

We assessed model fits to the data as per previous work (*29*), using the Kolmogorov-Smirnov (KS) statistic to quantify the distance between model and empirical network distributions of node degree, node clustering, node betweenness, and edge length distributions, with the largest such distance being taken as the final index of model fit (max(*KS*) i.e., the performance of a given model was assessed according to the property that it captured least accurately; Fig. 1D). To identify the best-fitting parameters for each model, we employed an optimization procedure that sampled 10,000 different parameter combinations, preferentially sampling from areas in the parameter landscape that produced the best fits (Fig. 1E; see Methods). Past models have evaluated model fitness based on within-sample performance, making it difficult to compare models with different complexity. Thus, to ensure that our results were not driven by overfitting, and that models with different numbers of free parameters could be compared fairly, we used a leave-one-out cross-validation procedure (note many different cross-validation schemes could be used, but we used leave-one-out as to limit computational burden associated with exhaustive parameter sweeps and model fitting). As depicted in Fig. 1F, this procedure comprised three steps: (1) for each individual’s data, we fitted models using the optimum parameter values obtained for the other 99 individuals from the initial sweep of 10,000 parameter combinations (with optimal parameters defined using max(*KS*)), and repeated this process 20 times to account for stochastic fluctuations in the models, yielding 99 × 20 model networks for each person; (2) we then took the average across the 20 runs, resulting in 99 mean fit estimates; (3) the average fit over these 99 models in the held-out individual was recorded; (4) Steps 1–3 were repeated so that each individual in the sample was held out once, resulting in a cross-validated fit-statistic, *F*_*CV*_, for each participant, with smaller values indicating a better fit.

The generative model forms connections probabilistically and one at a time according to a specific set of wiring rules. Under the traditional formulation of the cost-topology model proposed by previous studies (*28, 29*), the wiring rule can be written as

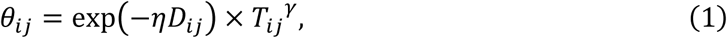

where *θ*_*ij*_ is connectivity score that is used to subsequently derive a probability of a connection forming between nodes *i* and *j* at a given time step (see Methods), *D*_*ij*_ is the distance between those nodes, *η* is a parameter controlling the scale of the distance decay, *T*_*ij*_ is some topological relationship between nodes between nodes *i* and *j*, and *γ* is a parameter controlling the scaling of the topological term. We note that prior work has often formulated the wiring cost term using a power-law, rather than exponential, distance-dependence but we use the exponential form here due to the abundant empirical evidence for such a dependence (*19, 20*), to allow a direct comparison to the widely studied EDR (*19, 20*), and because the scale-invariance of the power-law function means that any such models will preserve relative wiring costs under global changes in brain size, and will therefore be insensitive to the developmental changes in brain geometry introduced in our growth class of models. We followed the approach of Betzel and colleagues (*29*) and considered 12 different topological terms for *T*_*ij*_ that capture various aspects of degree, clustering, and connection homophily, the formal definitions of which are provided in Table 1.

**Table 1.**
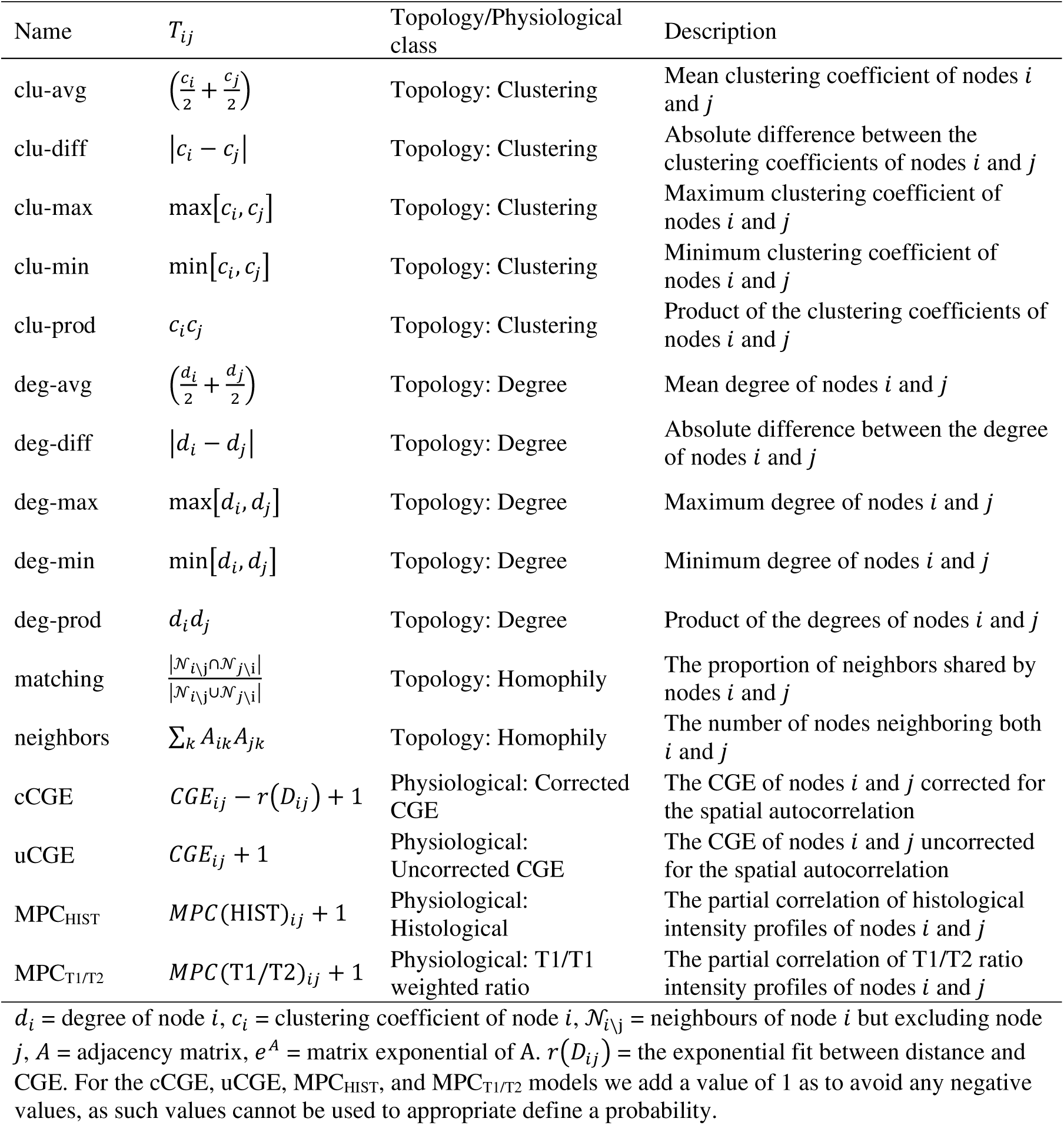
Definitions of topological terms used in the cost-topology generative models.

In the Methods and Supplementary Results, we show that the model formulation expressed in Eq. 1 can disproportionately penalize long-range connections and lead to ambiguous interpretation of parameter estimates due to a lack of independence between model parameters. We therefore derived a new formulation, given by

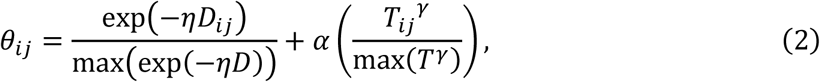

where *α* controls the contribution of the topological term and each term is normalized by its maximum value to ensure appropriate scaling of the distance and topological quantities. Practically, when estimating *θ*_*ij*_, high values of *α* assign more weight to the topological term; high values of *η* indicate a stronger distance penalty (i.e., shorter length scale of connectivity); and high values of *γ* control the non-linear scaling of the topological term, such that large values of *T*_*ij*_ exert a proportionally greater influence than smaller values. Models including the non-linear scaling of topology provided by *γ* fitted the data better than models that excluded *γ* (fig. S1, see supplementary text). Moreover, fig. S1C shows that the additive formulation of Eq. 2 more accurately captures putative trade-offs between cost and topology in connectome wiring, leads to more interpretable parameters estimates, and can fit our data better than the multiplicative formulation in Eq. 1, particularly with regard to capturing the empirical edge length distribution (fig. S2). Therefore, all results presented in the following sections use the basic formulation given in Eq. 2.

### 2.2 Accounting for developmental changes in cortical geometry

We first set out to determine whether incorporating developmental constraints into the generative models, by accounting for fetal changes in brain size and shape when estimating wiring costs, influences model performance. Fig. 2 shows how cortical geometry varies through time (Fig. 2A), along with variations in total surface area (Fig. 2B) and inter-regional distances (Fig. 2C). From 21 GA to 38 weeks GA, there is a 378% increase in total cortical surface area (Fig. 2B) and the distribution of inter-regional distances gradually becomes more skewed, such that an increasing number of regional pairs separated by longer anatomical distances emerges (Fig. 2C) and the maximum possible distance increases by 74%. In comparison to the adult brain, total cortical surface area observed at 38 weeks GA is 67% smaller and the maximum inter-regional distance is 41% shorter. Such large differences indicate that wiring costs estimates using adult brain geometry vary greatly from those that are likely to operate when inter-regional connections are being established.

**Fig. 2.**
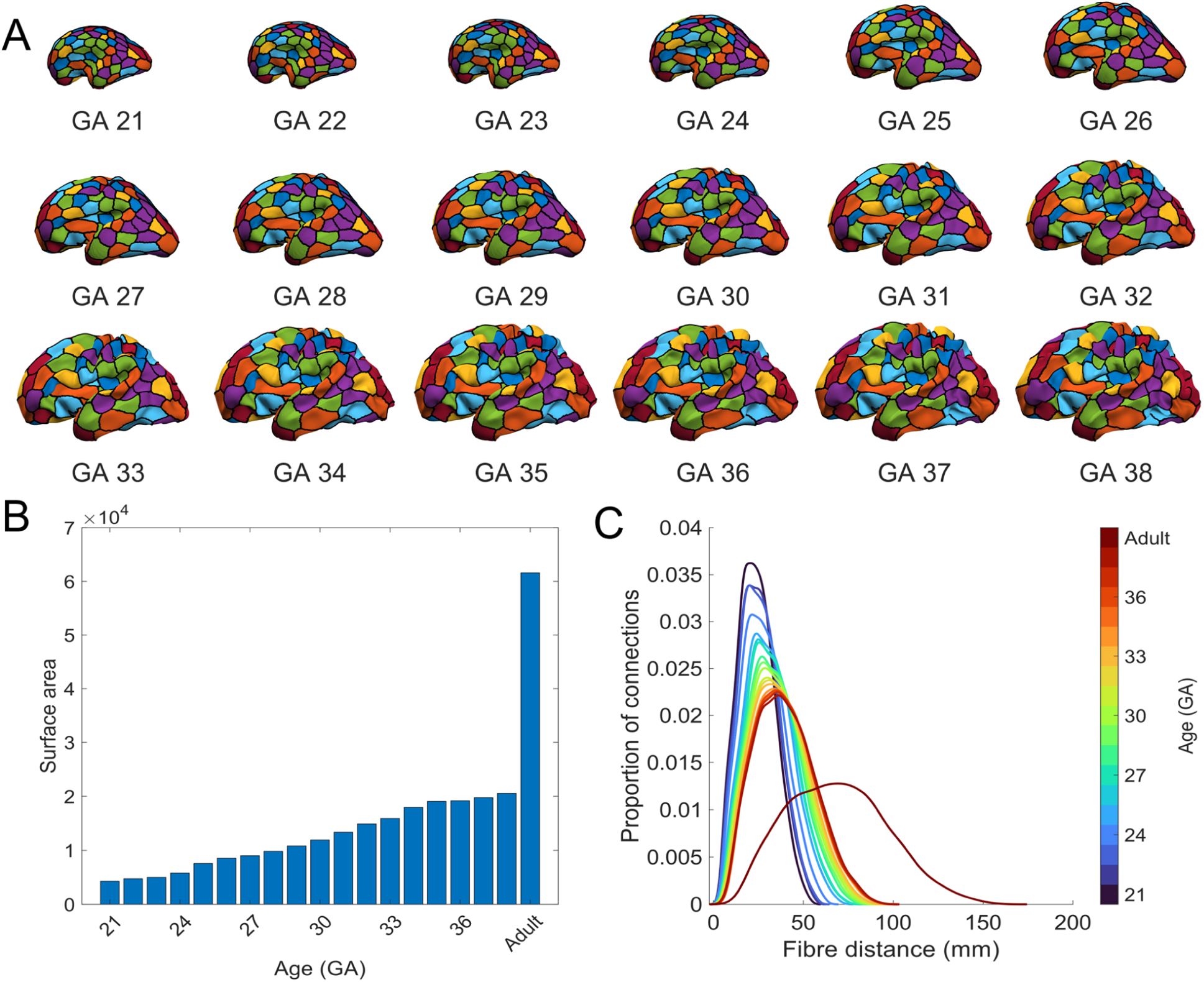
Surface area and fiber-distance distributions for the fetal surfaces. **(A)** The adult brain is warped into the shape of the fetal brain at each gestational age (GA) timepoint (see Fig. 1B) and allowing our node parcellation to be projected through developmental time. **(B)** Surface area estimated using the inner white/grey boundary of the left hemisphere of each fetal brain, with the adult included for comparison. **(C)** Kernel density plots of inter-nodal fiber distances between all nodes at each developmental time point.

To incorporate such developmental changes in brain size and shape into out models, we introduced a time-varying wiring cost to the basic formulation given by Eq. 2, yielding

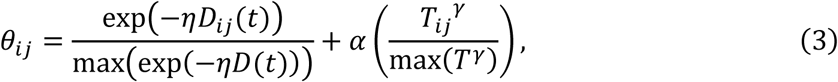

where *D*_*ij*_(*t*), which is the distance between nodes *i* and *j* at timepoint *t* (where *t* = 1 is 21 weeks GA, *t* = 2 is 22 weeks GA, and so on). For simplicity, we add connections at a uniform rate at each developmental time point, resulting in *E*/19 connections being added iteratively (i.e., one by one) according to the wiring costs given by any individual time point, with *E* corresponding to the number of edges in the empirical network, and 19 representing the number of developmental timepoints considered (18 fetal and one adult). Once the set number of edges is added at a given timepoint, the distances are updated to the next timepoint and the procedure repeats until *E* edges have been added (note that in the static model, all edges are added according to the same geometry).

For the static and growth trade-off models, we compared 12 different topological terms for *T*_*ij*_, as done previously (Betzel et al., 2016, see Table 1 for definitions) along with a purely spatial model based solely on the EDR. The cross-validated fit statistics for these models are shown in Fig.

3. We replicate prior work (*28, 29*) in showing that trade-off models generally out-perform the purely spatial EDR-based model, even when considering out-of-sample performance. While the spatial growth model yielded a small yet statistically significant improvement in mean *F*_*CV*_ relative to its static counterpart (growth 0.34 ± 0.03, static 0.35 ± 0.02, *p*_*FWER*_ < 0.001), this improvement was not sufficient to surpass the performance of the best-fitting trade-off models. Thus, even when developmental changes in brain geometry are considered, purely spatial models offer an incomplete account of the data. This result offers important confirmation of the hypothesis that, compared to a simple EDR process, the trade-off models more accurately capture the four key statistics of human connectome topology considered here.

As per past work (*29*), we found that the best-fitting model combines a distance penalty with a homophilic attachment rule based on the matching index (see Table 1). We also observed a significant (*p*_*FWER*_ < 0.001) performance advantage for the growth variant of this model over the static case, indicating that incorporating developmental constraints enhance the accuracy of this model, with an average improvement of ∼10% (i.e., the mean *F*_*CV*_ values for the best-fitting static and growth matching index models were 0.22 ± 0.01 and 0.20 ± 0.01, respectively). Performance differences between growth and static variants of the other cost-topology models were smaller.

As *F*_*CV*_ is derived from the worst KS statistic across four topological measures (node degree, node clustering, node betweenness, and edge length), we examined the extent to which the performance of each of these measures shaped the resulting *F*_*CV*_ value. For most models, the nodal clustering, nodal betweenness, and edge length distributions were the final determinant of the final *F*_*CV*_ value, indicating that these properties were the most difficult to capture (fig. S3). By comparison, the degree distribution was better captured (as indicated by only a small proportion of max(*KS*) values being determined by the KS statistic for degree). *F*_*CV*_ for the best-fitting matching model, in both static and growth cases, was more evenly shaped by the different topological properties. Notably, the growth matching model more accurately capture the empirical edge length distribution than its static counterpart, suggesting that improved performance of the growth model arose from a more accurate estimate of network wiring costs, as expected.

Comparison of model parameter estimates offers further insights into the relative behavior of the growth and static models. The optimal parameters for the two classes of models showed substantial differences; for instance, the growth matching model had best-fitting parameters of *η* = 0.35 ± 0.17, *γ* = 1.76 ± 0.44, and *α* = 2.05 ± 1.29, while the static model had *η* = 0.20 ± 0.15, *γ* = 1.30 ± 0.36, and *α* = 3.93 ± 2.15 (fig. S4). The higher *η* observed under the growth formulation suggests that an increased distance penalty is required for an optimum model fit compared to the static form. This effect likely arises because the length scales in the growth model are shorter than in the static variant, so the growth model requires a stronger distance penalty to match the adult edge-length distribution. Relative to the static variant, the growth matching model also required a weaker weighting and stronger non-linear scaling of the topology term, as indicated by the magnitudes of the *α* and *γ* parameters, respectively (fig. S4). This result suggests that connection probabilities were more heavily skewed towards node pairs with high matching index values in the growth variant. Given that the growth variant was also associated with a stronger distance penalty (higher *η*), the higher value of *γ* implies that topologically valuable connections are more likely to form during early stages of the growth model, when the distance penalty is weaker. Notably, for cost-topology models showing similar performance to the purely spatial model, *α* was approximately 0, consistent with the observation that topological rules did not improve model accuracy in these cases. These mechanistic interpretations of model parameters are only possible under our new model formulation (i.e., Eq. 2 and Eq. 3), as the classical formulation of Eq. 1 does not sufficiently separate the contributions of cost and topology (see Methods).

### 2.3 Physiologically-informed attachment rules

Our results indicate that cost–topology trade-off models offer a more accurate account of empirical human connectome topology than purely spatial, EDR-based models, and that homophilic attachment mechanisms informed by the matching index show the strongest performance. Our findings also indicate that incorporating developmental constraints into the matching model improves its accuracy. However, the topological homophily rule is an abstraction with no clear physiological mechanism. We next asked how the performance of this rule compares to models that incorporate alternative, physiologically grounded homophilic attachment mechanisms, such as those related to the architectonic type principle (*38, 41, 49, 50*). First, we estimated the microstructural profile covariance between pairs of regions using the Big Brain atlas, which is a Merker-stained 3D histological reconstruction of a post mortem adult human brain (*51*). This measure, which we term MPC_HIST,_ quantifies inter-regional similarity in estimates of cell size and density through the cortical depth (*52*). Second, we estimated microstructural profile covariance derived from the ratio of T1-weighted to T2-weighted signal estimated from in vivo MRI (MPC_T1/T2_), which is often used as a proxy for intracortical myeloarchitecture (*53*). Finally, given the reported link between coupled gene expression and neuronal connectivity (*5, 23*), we also evaluated measures inter-regional transcriptional coupling, quantified as correlated patterns of expression measured across 1634 brain-expressed genes (*54*) using data from the Allen Human Brain Atlas (AHBA; (*55, 56*)). We term this measure correlated gene expression (CGE) (*5, 23*). Further details are provided in the Methods.

For each of these three physiologically-informed models, we evaluated model performance with or without a wiring-cost term in the model. Models without the wiring cost take the form *θ*_*ij*_ = *T*_*ij*_ ^*γ*^, where *T*_*ij*_ is either MPC_HIST_, MPC_T1/T2_, or CGE. Models with wiring cost include *D*_*ij*_, as per in Eq. 2 or use *D*_*ij*_(*t*) as in Eq. 3 when considering growth variants. Note that growth variants can only be estimated for models that include wiring costs. For CGE-constrained models, we examined variants that used either raw CGE values (uCGE) or values corrected for the well-known distance-dependence of the coupling estimates (cCGE) (*23, 56*). The relationship between MPC and distance is more regionally variable and a bulk distance correction unevenly impacts certain areas (*57*) and so we only consider raw estimates of these quantities (see Methods).

First, we examined whether any of the single-parameter physiological models (MPC_HIST_, MPC_T1/T2_, cCGE, or uCGE) could outperform the classical spatial and/or cost-topology models. As shown in Fig. 4, all but the cCGE outperformed the spatial model (*p*_*FWER*_ < 0.001), but only the uCGE model (*F*_*CV*_ = 0.18 ± 0.02) outperformed the matching index model (static: *F*_*CV*_ = 0.21 ± 0.01; growth: *F*_*CV*_ = 0.20 ± 0.01). Adding a distance to term to the physiological models improved the performance of the MPC_HIST_ and MPC_T1/T2_ models, and growth variants were associated with slight performance advantages for all but the spatial+ MPC_T1/T2_ model. Critically however, none of these models surpassed the accuracy of the uCGE model. Moreover, combining uCGE with a wiring cost term offer only a minor 3% performance gain (growth spatial+uCGE: *F*_*CV*_ = 0.182 ± 0.019; uCGE: *F*_*CV*_ = 0.188 ± 0.017), suggesting that the single-parameter uCGE model offers a parsimonious account of the data.

**Fig. 3.**
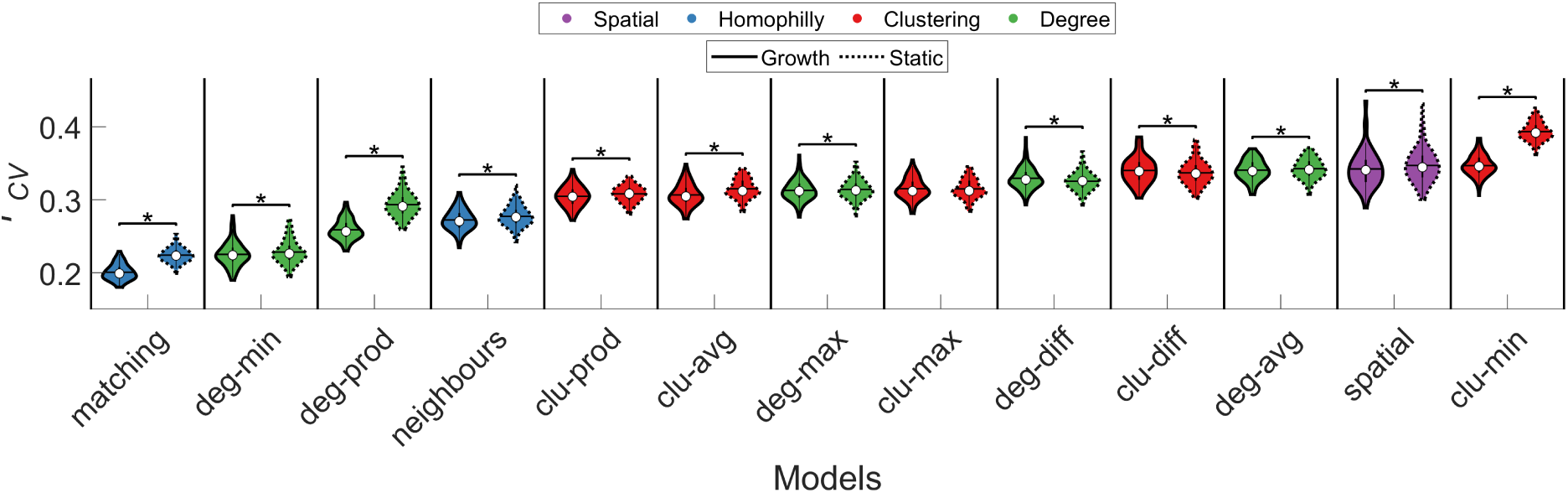
Model performance for static and growth variants of spatial and cost-topology trade-off models. Each violin plot shows the distribution of cross-validated *F*_*CV*_ values for static and growth additive models across subjects. The color of each violin plot indicates the topology metric used in the model: homophily is shown in blue, clustering (clu) in red, degree (deg) in green, and spatial in purple. The white circle indicates the median of each distribution, while the horizontal black line indicates the mean. The matching growth model achieved the best performance. * *p* < 0.05 Bonferroni corrected (325 tests between all 26 models), Wilcoxon signed-rank test.

**Fig. 4.**
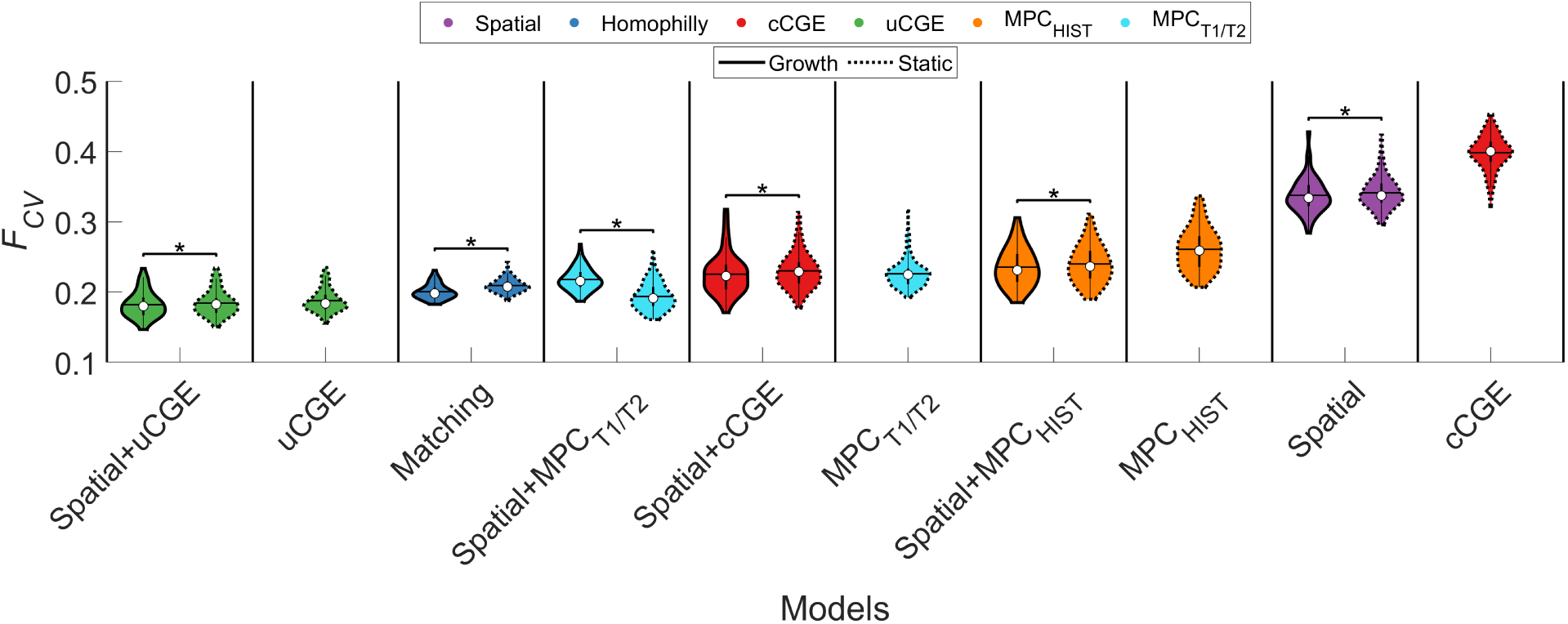
Model performance of physiologically-informed models. Each violin plots shows the *F*_*CV*_ values different models. The color of each violin plot indicates the type of model: matching is shown in blue, cCGE in red, uCGE in green, MPC_HIST_ in orange, MPC_T1/T2_ in cyan, and spatial in purple. The white circle indicates the median of each distribution, and the horizontal black line indicates the mean. uCGE models achieved the best fit. Note that uCGE, cCGE, MPC_HIST_, MPC_T1/T2_ models do not have a growth variant as they do not have an independent distance term. * *p* < 0.05 Bonferroni corrected (120 tests between all 16 models), Wilcoxon signed-rank test.

The lack of improvement observed with the addition of a distance penalty to the uCGE model is likely due to the approximately exponential distance-dependence that is already present in uCGE values (*23, 56*). Indeed, as noted above, the cCGE model, which explicitly removes the intrinsic spatial dependence of CGE, was the worst performing model. This result, together with the strong performance advantage of the uCGE model over the purely spatial model, indicates that it is the specific spatial patterning of CGE, beyond a simple EDR-based distance dependence, that is particularly informative about connectome topology. The only other physiologically informed model to out-perform the matching index model was the static variant of the spatial+ MPC_T1/T2_ (*F*_*CV*_ = 0.19 ± 0.025, *p*_*FWER*_ < 0.001). All other physiologically informed models showed either comparable or slightly worse performance than the matching-index model.

Evaluation of model fits to the specific topological properties used in the fitting procedure indicated that the physiological models (excluding cCGE, spatial+cCGE, and MPC_T1/T2_) more accurately captured the edge length distribution than the topological models, for which edge lengths were fitted least accurately (fig. S5). For the physiological models, betweenness and clustering were the two properties that were least accurately reproduced.

### 2.4 Modeling topographical properties of the human connectome

Our findings indicate that physiologically-informed homophilic attachment mechanisms, and particularly those constrained by inter-regional transcriptional coupling, can reproduce key topological properties of the human connectome better than wiring rules based on topological homophily. We next investigated whether these models can also reproduce the way in which these properties are spatially embedded; i.e., the topography of the connectome. To this end, we focused on the performance of the uCGE models, matching index, and spatial models in in reproducing the spatial topography of regional clustering, betweenness, degree, and mean connection distance; that is, the spatial characteristics of the four topological properties used to fit the models to the data. We quantified model performance in capturing topographical properties as the Spearman correlation between the best-fitting model and empirical node sequences for each of these properties. Note that only the topological (i.e., statistical distributions), and not topographical (i.e., node/edge sequences), properties were used to optimize model parameters.

We found that while the uCGE and matching models closely capture the statistical distributions of topological features, all models generally show poor performance in capturing topographical properties, with the average correlation across subjects never exceeding 0.21 (Fig. 5). Despite this generally modest performance, the uCGE and spatial+uCGE models showed better performance across nearly all topographical properties.

**Fig. 5.**
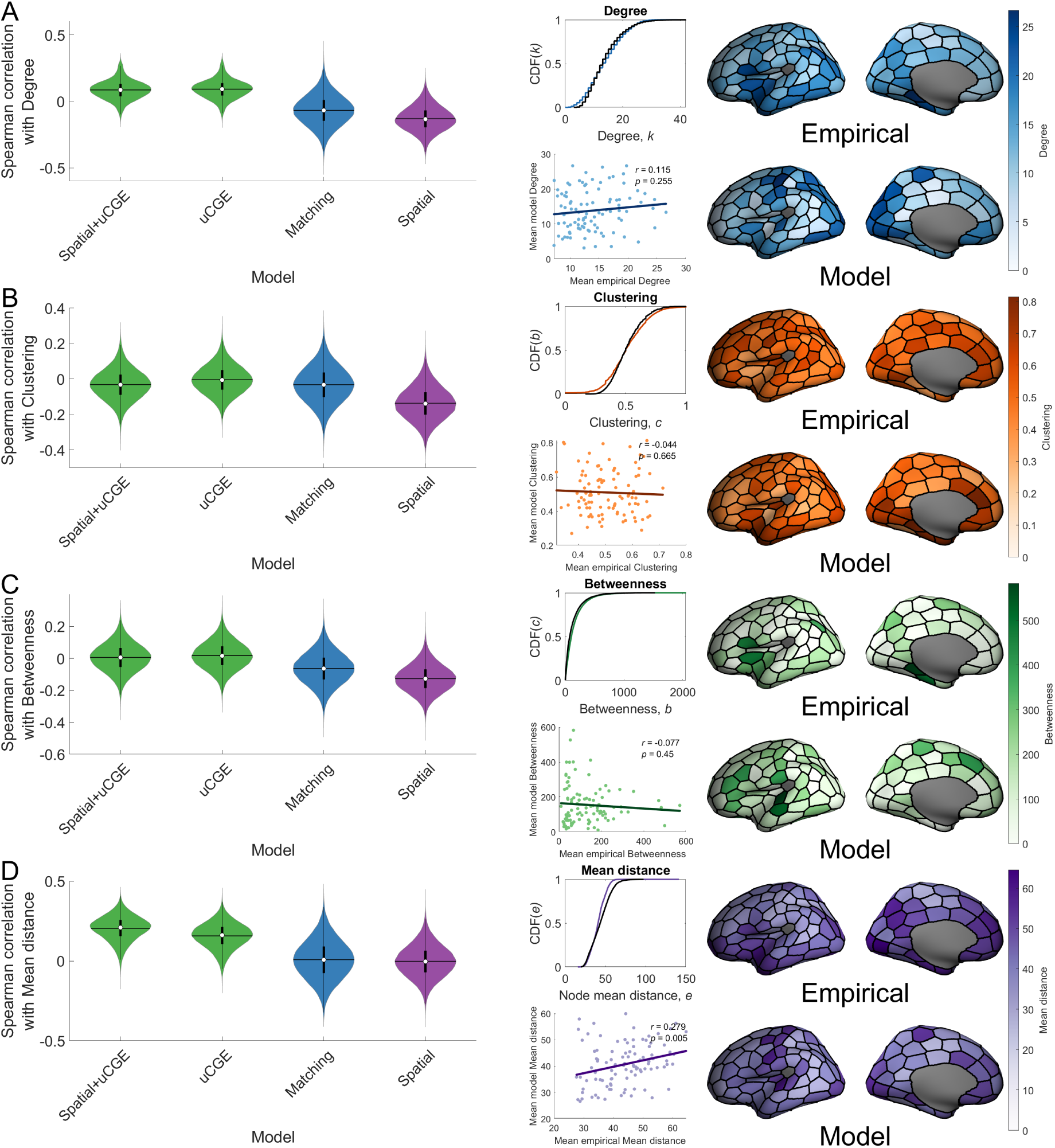
Model performance in capturing connectome topography. For each of the network measures that were used to evaluate model performance, we show violin plots of the spearman correlation between the empirical and data for selected models for a given property of the model fit function. For the spatial+uCGE model, we additionally show, for each network measure, the average CDF of the model data (colored line) as compared to the empirical data (black line); a scatter plot of average nodal model values against average nodal empirical values; and a projection of these across-subject average nodal measures onto the cortical surface. **(A)** Node degree spatial arrangement. **(B)** Node clustering spatial arrangement. **(C)** Node betweenness spatial arrangement. **(D)** Mean nodal distance spatial arrangement.

The models considered in Fig. 5 were optimized to fit topological properties. To investigate in more detail whether the models can accurately reproduce topographical properties regardless of fits to topological distributions, we evaluated the maximum the Spearman correlation between empirical and model node degree obtained across all parameter combinations evaluated in our model fitting procedure. We found that no such correlations ever exceeded 0.49, with median correlations across the models ranging between −0.17 and 0.13 (fig. S6), further suggesting that current generative models have a limited capacity for reproducing topographical properties of the human connectome.

## 3 Discussion

In this work, we introduce a new formalism for accurately capturing how cost-value trade-offs might shape brain network wiring, and combine this new model with a framework for incorporating physiological constraints and developmental changes in brain size and shape. Using a cross-validated model evaluation procedure that accounts for variations in model complexity, we show that developmentally informed growth models fit the data better than models assuming fixed wiring costs through development. As per prior work (*5, 28–32*), we confirm that cost-topology trade-off models perform better than purely spatial models, but also show that physiologically-constrained models, particularly those in which the probability of forming a connection between two regions is influenced by their level of transcriptional coupling, offer a more accurate and parsimonious account of connectome topology. While physiological models did show better reproduction of empirical topographical properties than cost-topology models, all models weakly captured the way in which the data are spatially embedded. Collectively, our findings suggest that a simple, single-parameter generative model with a homophilic attachment mechanism based on transcriptional coupling offers the most parsimonious account of connectome topology, and that additional constraints may be required to accurately model topographical properties of human brain networks.

### Parsing the effects of space, topology, and physiology on connectome wiring

The EDR has been proposed as a fundamental constraint on neuronal connectivity, having been used to explain edge-length distributions and the presence of particular kinds of cliques and motifs in the connectomes of the mouse and macaque (*19, 20*). The rule implies that stochastic processes subject to a distance-dependence are sufficient to explain connectome topology. Past work directly comparing EDR-based models to cost-topology trade-off models has found that the latter class fit empirical macroscale human connectome data better (*5, 28–30, 32*), but these studies did not account for differences in model complexity (see Methods). Our cross-validated fitting procedure allowed fair model comparison and confirmed the superiority of the trade-off models. Moreover, we showed that incorporating developmental changes in brain geometry still resulted in superior performance for trade-off compared to spatial models, indicating that a past reliance on using adult estimates of wiring cost has not artificially limited the performance of models based solely on EDR-like processes. These findings are in line with Cajal’s (*7*) hypothesis that an interplay between wiring cost and functional value shapes brain network wiring.

Of the trade-off models considered here, those relying on homophilic attachment guided by the matching index performed better than models based on properties of node clustering or degree, consistent with prior work (*28–30, 32*). However, while this form of topological homophily may be plausibly linked to a Hebbian-like plasticity process (*33*), the precise mapping between topological terms such as the matching index and the physiological processes that sculpt neuronal connectivity remains unclear. We therefore investigated an alternative class of homophilic attachment models in which inter-regional homophily was informed by physiology rather than topology and showed that such models often perform better than the matching index model.

In a general sense, all three physiologically-constrained models considered here––CGE, MPC_HIST_, and MPC_T1/T2_––offer different ways of testing the architectonic type principle, or structural model of neuronal connectivity, which states that regions with more similar cytoarchitecture and laminar organization and more likely be connected with each other (*40, 41, 49, 58*). MPC_HIST_ and MPC_T1/T2_ represent more direct measures of cytoarchitectonic similarity, quantifying cortical depth-dependent variations in cell size/density and myelin content, respectively. CGE offers an arguably less direct, although perhaps related measure of microstructural similarity. Gene-expression measures in the AHBA are obtained through bulk tissue microarray, and the resulting expression values will also be influenced by regional variations in cellular architecture, in addition to other factors influencing regional transcriptional activity, although the specific contributions to CGE made by cytoarchitectonic or other aspects of transcriptional similarity remain unclear.

The superior performance of both CGE and MPC_T1/T2_ compared to MPC_HIST_ models may indicate that inter-regional coupling of factors related to myeloarchitecture may be more closely linked to connectivity than similarity in neuronal organization, in light of evidence that the T1/T2 ratio track intracortical myelin (*53*) and that oligodendrocyte-related genes contribute to variations in CGE that are linked to inter-regional connectivity (*5*) Notably, MPC_T1/T2_ has a lower resolution than MPC_HIST_, and is thus more sensitive to the skewness of the intensity profiles, which varies along a sensory-fugal axis (*57*). As such, MPC_T1/T2_ more closely corresponds to the hierarchical sensory-fugal axis, which is an established organizing principle of neuronal connectivity (*59*). Additionally, the BigBrain atlas from which MPC_HIST_ estimates were derived was constructed using only a single brain and the challenges of data reconstruction and lack of averaging across individuals may result in somewhat noisier measures.

The spatial+uCGE model showed the lowest average *F*_*CV*_, closely followed by the single parameter uCGE model. The strong performance of the single-parameter uCGE model suggests that a non-linear (power-law) scaling of CGE values to favor connections between regions with positive CGE (see Methods) provides a parsimonious model of macroscale connectome topology, with little additional benefit from the inclusion of a term for connection wiring costs. Although the uCGE values show a strong and approximately exponential distance-dependence (*23, 56*), this dependence alone cannot account for the strong performance of the uCGE model, given that the spatial model performed so poorly. Rather, it is the specific spatial patterning of CGE values that is likely to be important in shaping connectome topology. This conclusion is further supported by the poor performance of the spatial+cCGE model, which replaces the intrinsic distance-dependence of the CGE values with a fitted exponential wiring-cost penalty. Thus, while a bulk exponential trend can approximate the distance-dependence of CGE, fluctuations around this trend may pay a central role in shaping inter-regional connectivity. In this sense, both spatial and physiological constraints may be more relevant to understanding connectome wiring than abstract topological rules.

### Accounting for developmental changes in brain size and shape

In general, growth-based model variants yielded small, yet significant, performance advantages over their static counterparts when considering the best-fitting models. This result suggests that developmental changes in cortical geometry may not play a substantial role in shaping connectome topology. Optimal values of *η*, which define the distance penalty imposed in the model, were larger in the growth models than the static model, indicating that an increased distance penalty was required for the growth models, on average. This increased penalty counteracts the potential benefit of shorter distances at earlier timepoints. Since the position of nodes, and thus the relative distances between them, did not change drastically from one timepoint to another (except for between the 38 gestational week and adult brain timepoints), it is likely that the models fitted the *η* parameter to the average distance across all these timepoints. Since the distances in the fetal brains largely represent scaled-down adult distances, this behavior will limit potential performance differences between growth and static model variants. One way around this limitation is to fit a distinct value of *η* at each timepoint. Future work could also look to vary the rate at which connections are added at different time points, however, these changes come at the cost of an explosion in model complexity. We opted for the simplest approach and fixed *η* across time, but our basic framework could be adapted to investigate these more nuanced influences in future work.

Our growth models were implemented such that any node at any given time point could form a connection. This is known as tautochronous (or parallel) growth. Simulations have suggested that heterochronous (or serial) growth, in which there is a prescribed order to which nodes can form connections, may offer a more realistic model (*37, 49, 60, 61*). Heterochronous growth is thought to play an important role in shaping the relationship between cytoarchitectonic similarity and connectivity (*58*) and may facilitate the formation of long-range connections when combined with spatial changes in brain geometry. Our modeling framework can be extended to consider heterochronous growth by adding connections at different rates or times for different brain regions. Developing principled ways of parametrizing such growth processes will be an important extension of the current work.

### Modeling topographical properties of the connectome

Initial generative modeling studies only evaluated model performance with respect to a small set of hand-picked topological properties (*28, 29*), thought to be characteristic of the human connectome. Only recently have topographical properties been considered (*5, 30–32*). Recent studies have shown the importance of the spatial location of high-degree hub regions (*42*), and indicated that classical cost-topology trade-off models cannot accurately capture the spatial arrangement of nodal degree (*31*) (however see (*30*) as well as limitations below), while incorporating transcriptional information can improve accuracy (*5*). These findings motivated us to consider the extent to which our models could reproduce the topographical properties of degree, clustering, betweenness and mean connection distance. We replicated prior results indicating that physiologically constrained models, particularly those using CGE estimates, more accurately captured diverse aspects of network topography (*5*). However, even the best-fitting model achieved only moderate success, with the highest spatial correlation in the spatial+uCGE model (across all four properties) being *ρ* = 0.19. It is thus possible that a combination of transcriptional constraints and heterochronous growth may be necessary to accurately capture both topological and topographical properties of the human connectome. This combination may result from developmental variations in transcriptional profiles guiding axons to their targets. The construction of anatomically comprehensive gene-expression atlases through different stages of prenatal development (*62*) would help to test this hypothesis.

#### 3.1 Limitations

The reference cortical surfaces we used at each fetal timepoint were obtained from different fetuses and using differing numbers of scans, introducing variability in cortical shape and size between gestational time points (*45, 46*). This variability means that our surface model does not smoothly develop from one timepoint to another, as would be expected in an actual brain. Nonetheless, we expect that the fiber distances estimated using these geometries should not vary dramatically, and that for present purposes they represent a reasonable first approximation of developmental changes in geometry.

Our growth model added connections one at a time, but connections are likely to form contemporaneously in the developing brain. Connections are added sequentially in the model so that the topology of each edge can be recalculated at each iteration but future extensions may consider sampling multiple edges at any given time. Moreover, we added an approximately equal number of connections at each developmental timepoint, but more complex temporally and spatially varying schemes for connection formation are possible. Future work could also consider modeling physical processes of connectivity in geometric space, like specific rules for axonal guidance or neurodevelopmental gradients, in order to provide a stronger mechanistic account for how connections form (e.g., see (*25, 61*)).

In line with previous studies (*28–30, 32*), we only examined the ability of generative models to predict binary network topology. However in empirical connectomes, edges have weights which span several orders of magnitude (*63*). Since physiological factors which wire the brain are linked to variation in connectivity strength (*58*), future work should look to adapting generative models to produce weighted networks. Extending the framework developed here to capture weighted network properties represents an important extension of our work.

Finally, while numerous studies have used generative models, they have been fitted to connectomes generated using different preprocessing pipelines, thresholding methods, parcellations, one or both hemispheres, and many other methodological variations. As the topology of brain networks can differ based on how the data were processed (*64–66*), it is possible that such variations may influence model performance. For example, studies using higher-resolution parcellations comprising ≥ 100 nodes have encountered difficulty in replicating the spatial embedding of network hubs (*5, 32*), whereas one study using a lower-resolution parcellation of 68 nodes performed better in relation to node degree (*30*). Moreover, studies using probabilistic rather than deterministic tractography have identified alternative cost-topology models to the matching index model as offering the best fit to empirical data (*5*). A better understanding of how model performance depends on data preprocessing will be essential if the field is to converge on a parsimonious consensus model.

#### 3.2 Conclusions

We advance a framework for modeling the influence of cost-topology trade-offs in brain network development that allows fair comparison between models of different complexity, which captures developmental changes in brain geometry, and which can be used to incorporate additional physiological constraints. We show that simple, physiologically constrained models offer more accurate accounts of human brain topology than models relying on more abstract topological rules of the connectome, but that all generative models have trouble replicating topographic properties. Taken together, our findings suggest that geometric constraints and developmental variations in regional transcriptional profiles may conspire to shape both the complex topological properties and specific spatial embedding of macroscale brain network architecture.

## 4 Methods

### 4.1 Data

We used data from the Human Connectome Project (HCP), randomly selecting images for 100 unrelated participants (49 female, age mean ± standard deviation: 28.79 ± 3.67). Data were acquired on a customized Siemens 3T Connectome Skyra scanner at Washington University in St Louis, Missouri, USA using a multi-shell protocol for the DWI with the following parameters: 1.25 mm^3^ voxel size, TR = 5520 ms, TE = 89.5 ms, FOV of 210×180 mm, 270 directions with *b* = 1000, 2000, 3000 s/mm^2^ (90 per *b* value), and 18 *b* = 0 volumes. Structural T1-weighted data were acquired with 0.7 mm^3^ voxels, TR = 2400 ms, TE = 2.14 ms, FOV of 224×224 mm (*44, 67*). A total of 100 participants were used due to the computational burden of running multiple different models for each participants network.

### 4.2 Connectome mapping

The HCP data were processed according to the HCP minimal preprocessing pipeline, which included normalization of mean *b* = 0 images across diffusion acquisitions and correction for EPI susceptibility and signal outliers, eddy-current-induced distortions, slice dropouts, gradient-non-linearities and subject motion. The details of this pipeline are provided in more detail elsewhere (*44, 68*). T1-weighted data were corrected for gradient and readout distortions prior to being processed with FreeSurfer (*44*).

To define network nodes, we parcellated the brain into 100 regions of approximately equal size. This parcellation was generated by randomly subdividing the fsaverage template surface. We only considered cortical regions for the parcellation as our approach to registering and aligning fetal brains did not extend to non-cortical areas. The parcellation was then registered from the template surface to the surface of each individual subject using a spherical registration procedure implemented in FreeSurfer (*69*), where it was converted to a volumetric image for subsequent network generation. We focus here only on the left cerebral hemisphere when performing generative modeling to follow past work (*5, 28, 29, 32*) and to reduce computational burden. While there are many different ways of parcellating human brain imaging data, we took a pragmatic view, requiring that a) parcels were of approximately equal size, since variations in regional size can affect many nodal properties, such as node degree; and b) the resulting networks were small enough that they could be modelled with sufficient computational efficiency, due to the large number of model iterations that we ran. Examining how model performance varies across different parcellations and data processing strategies is an important extension of the current work.

Deterministic tractography was performed using the Fiber Assignment by Continuous Tractography (FACT) algorithm (*70, 71*) as implemented in MRtrix3 (*72*). The algorithm propagates streamlines in the direction of the most collinear fiber orientation estimated within the voxel in which the streamline vertex resides. We defined one fiber orientation in each voxel by estimating the diffusion tensor using iteratively reweighted linear least squares (*73*) and then calculating the primary eigenvector of water diffusion. A total of 10 million streamlines were generated for tractography, with a maximum curvature of 45° per step. Streamline seeds were preferentially selected from areas where streamline density was under-estimated with respect to fiber density estimates from the diffusion model (*74*). Anatomically Constrained Tractography was used to further improve the biological accuracy of streamlines (*75*). To create a structural connectivity matrix, streamlines were assigned to each of the closest regions in the parcellation within a 5mm radius of the streamline endpoints (*76*), yielding an undirected 100 × 100 binary connectivity matrix (density 0.14 ± 0.01).

### 4.3 Mapping developmental changes in cortical geometry

To estimate developmental changes in cortical size and shape we obtained MRI scans from a public database of fetal MRIs (*45, 46*) acquired from 21-38 weeks GA. Most evidence suggests that the majority of axons form in this period, with nearly all inter-regional connections being formed by birth (*77, 78*). We therefore restricted our focus to this developmental window, but note that our framework can be flexibly extended to include estimates of post-natal cortical geometry.

The fetal scans are released as group average templates of scans available at each timepoint. For each brain, we manually segmented the T1 weighted images using ITK-snap (*79*) to label the white-matter mask, as existing automated segmentation algorithms suffer from poor accuracy due to the inherently poor tissue contrast in fetal images. Surfaces were constructed from the white-matter mask and smoothed using a heat kernel smoothing algorithm (*80*) and up-sampled by a factor of four (using four-split spline interpolation) to ensure an adequate number of vertices were available to perform the surface-based registration. For each extracted surface, we estimated maps of sulcal depth and projected the surface to a sphere using FreeSurfer (version 5.3).

To match these prenatal surfaces to the adult cortical surfaces, we used the Multimodal Surface Matching (MSM) algorithm (*47*). MSM matches an input and reference surface via their spherical projections. The algorithm warps the vertices on the input surface to maximize the similarity between a specified feature (in the case of this study, sulcal depth) of the two surface meshes while also minimizing the extent of this distortion (*47*). Additionally, higher-order clique reduction was used to improve surface regularization (*48*). This approach was used to register each fetal surface to the MNI 305 average surface template (fsaverage). More specifically, to prevent any bias due to the direction of registration (*81*), we took the average result of the registration of the fetal to the adult and the adult to the fetal surfaces (i.e., the mean coordinates of corresponding pairs of vertices in the two registrations was taken). This procedure allowed us to register the adult parcellation to each fetal surface, thus enabling us to track the spatial location of each network node through development.

To ensure the accuracy of our cortical surface model, we calculated cortical surface area and inter-regional fiber-distance distributions at each timepoint. The total surface area of each fetal brain showed an approximately linear increase over time (Fig. 2B). These values and trends are similar to those found in other studies reporting surface area changes in this developmental period (*82, 83*). Changes in estimated fiber distance for all possible pairs of brain regions are shown in Fig. 2C and confirm that distances between nodes gradually increase throughout development.

### 4.4 Estimating wiring costs

A true estimate of neuronal wiring costs requires a full consideration of the metabolic resources required to form and maintain connections between neurons. The data required for such consideration at the level of the entire brain are currently unavailable. As a proxy, most investigators use the physical distance of a connection to index wiring cost, under the assumption that longer connections require greater cellular material and physical space, and thus consume greater metabolic resources (*2*). Most studies in the field have approximated connection distances using the Euclidean distance between brain regions (*5, 28, 29, 32*). This approach can under-estimate actual fiber distances, as Euclidean distances do not account for the complex geometry of the cortex and do not track actual fiber trajectories through the white-matter volume. This is an especially pertinent consideration when assessing developmental changes in wiring costs, as the formation of sulci and gyri represents a prominent geometric change that may significantly alter distances between cortical areas over time. We thus aimed to more closely estimate actual fiber distances in our analysis by approximating the physical paths between brain regions that pass through the white-matter volume.

While it is straightforward to measure the length of reconstructed tracts, our models require wiring-cost estimates for all possible connections, including those that have not been empirically constructed, across different developmental time points. We therefore used the following procedure to generate such estimates. First, we downsampled the cortical surface using MATLAB’s *reducepatch* command (fig. S7A) so that only 15% of the original number of vertices remain (fig. S7B). This step preserves the shape of the brain but ensures efficient computation. Second, for each vertex in the downsampled surface model, we found a corresponding point located 0.1 mm interior and perpendicular to the surface (this step avoids precision issues that can occur in subsequent steps; fig. S7C). Third, we used ray tracing to draw a line segment between every pair of subsurface points and assess if this segment intersects the original surface mesh (figs. S7D-E). Fourth, a direct vertex connection matrix, *L*, was defined where each element *L*_*uv*_ indicated the Euclidean distance between vertices *u* and *v* if a line segment that did not intersect the surface could be drawn between their corresponding subsurface points; otherwise *L*_*uv*_ = 0. Finally, Dijkstra’s algorithm was run on the *L* matrix to find the shortest distance to connect each vertex through the interior of the surface (fig. S7F). We then took the average distance between all pairs of vertices in ROIs *i* and *j* to estimate the minimum possible fibre distance between each region/node. Note that the actual distances of fibers between regions are likely larger as they will be affected by factors such as fiber volume, ventricles, and subcortical structures. Our approach nonetheless offers a more accurate approximation of actual fiber distances than Euclidean distances.

### 4.5 Transcriptomic data

We constrained our generative models using transcriptomic data from the Allen Human Brain Atlas, which comprises 3702 spatially distinct tissue samples taken from six neurotypical postmortem adult brains (*55*). Across these brains, samples from 58,692 probes, distributed across cortical, subcortical, brainstem and cerebellar regions, quantify the transcriptional activity of 20,737 genes. As only two of the brains in the dataset sampled the right hemisphere, we exclusively focused our analysis on the left cortex. The preprocessing procedures applied to this data are described in detail elsewhere (*5, 56*). Briefly, genes were assigned to probes using the Re-Annotator toolbox, resulting in 45,821 probes and a corresponding 20,232 genes being selected. Samples annotated to the brainstem and cerebellum were removed, then intensity-based filtering was used to exclude probes which did not exceed background noise in more than 50% of samples. From the remaining 31,977 probes and 15,746 genes that survived filtering, a representative probe for each gene was selected based on the highest correlation to RNA sequencing data in two of the six brains. Samples were classified based on their hemisphere (left/right) and structural assignment (cortex/subcortex) assigned to regions of the 100 node parcellation by a) generating a parcellation for each doner specific brain and b) assigning samples to the closest region that matched their hemisphere and structural assignment within 2 mm of the parcellation voxels. Any samples assigned to subcortical or left regions were removed. Gene-expression measures were normalized within each region by first applying a scaled robust sigmoid normalization for every sample across genes, and then for every gene across samples. This normalization yields estimates of the relative expression of every gene across regions when controlling for donor-specific differences in gene expression. By averaging the normalized expression measures in each region across donor brains, we obtained a matrix of expression values for 10,027 genes in 99 regions (one region was removed as no samples could be assigned to it). We focused on a subset of 1634 genes that have previously been identified as expressed in human brain tissue (*54*). To quantify CGE for each pair of regions, we estimated the Pearson correlation between the normalized expression measures of the 1634 genes available after pre-processing.

The gene expression measures we used were obtained in adult specimens and the resulting CGE estimates may not directly reflect expression in the developing brain. While many genes show neotenous expression patterns (*84*) many others show highly variable expression patterns though development (*85*). Present transcriptional atlases of the developing human brain lack the anatomical coverage to allow estimation of whole-brain CGE profiles (*62*), although analyses in mouse indicate a predictable scaling rule in the distance dependence of CGE throughout development (*86*). As the coverage of these atlases improves, developmentally varying CGE estimates could be readily incorporated into the growth class of models introduced here.

It is well-documented that the level of transcriptional coupling between two regions declines as an approximately exponential function of the distance between them (*22, 23, 56, 86*). This spatial autocorrelation is physiologically meaningful and may be fundamental to the relationship between gene expression and brain connectivity (*23*). However, it can also be informative to disentangle CGE estimates from their distance dependence. We therefore incorporated two types of CGE estimates into our generative models: (1) CGE estimates corrected for their intrinsic spatial autocorrelation (cCGE); and (2) uncorrected, raw CGE estimates (uCGE). The cCGE estimates were obtained by fitting an exponential function with the form 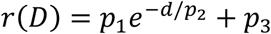, where *p*_1_ = 1.12, *p*_2_ = 0.023, and *p*_3_ = −0.27 (distances along the cortical surface were used to calculate this function, as done in (*56*)). The residuals of this fit were used in the modeling as cCGE values, which for each pair of regions was defined as *cCGE*_*ij*_ = *CGE*_*ij*_ − *r*(*D*_*ij*_). The cCGE and uCGE estimates were remapped to the positive range by adding a constant, *c* = 1, to all values, to ensure that our models did not return negative connection probabilities. As such, the scaling exponent *γ* applied to CGE estimates in our models serves to strongly weight pairs of regions with positive compared to negative CGE values.

### 4.5 Microstructural profile data

In addition to transcriptional coupling, we investigated two measures of microstructural profile covariance (MPC) between regions. One used histological data from the BigBrain atlas (*51*), a Merker-stained 3D volumetric histological reconstruction of a human brain (MPC_HIST_). Following Paquola et al. (*52, 57*), we constructed 50 equivolumetric surfaces between the white and pial surface boundaries and then sampled the intensity values along these surfaces at each vertex. MPC_HIST_ was then obtained by taking the partial correlation of regional mean intensity profiles whilst controlling for the cortex-wide mean intensity profile (as with the CGE models, MPC results are transformed into the range 0-2). MPC_HIST_ can thus be interpreted as a measure of inter-regional similarity in variations of cell density and size through the cortical depth.

To estimate MPC_T1/T2,_ we applied a similar approach to the T1/T2-weighted ratio obtained with in vivo MRI in an independent sample of 197 unrelated healthy adults from the Human Connectome Project. For each subject, 12 equivolumetric surfaces between the inner and outer cortical surfaces were constructed and used to sample T1/T2 values across each vertex (*52*). MPC was then calculated as with the BigBrain atlas, and an average was taken across all subjects to obtain a single MPC_T1/T2_ data matrix. To the extent that the T1/T2-weighted ratio indexes intracortical myelin (*53*), MPC_T1/T2_ can be interpreted as an indirect measure of inter-regional similarity in myeloarchitectonic variations through the cortical depth. Both MPC_HIST_ and MPC_T1/T2_ show subtle distance-related trends that are not easily accommodated with bulk corrections. We therefore consider only raw, uncorrected estimates in our analyses.

### 4.6 Generative modeling

#### Basic model characteristics

Several different types of generative models for connectomes have been proposed (see Betzel & Bassett, 2017 for a review). We focus here on the cost-topology trade-off model, as defined in Eq. 1, which has been extensively studied in the content of human brain networks (*5, 28–30, 32*). The model defines a relation between wiring cost (e.g., *D*_*ij*_) and topology (e.g., *T*_*ij*_) that influences the probability of forming an edge between two nodes *i* and *j*. Edges are added one at a time to the network, with the topology value being recalculated at each iteration and connection scores updated accordingly. The model is iterated until the number of edges in the synthetic networks matches the empirical data.

Following prior work (*28, 29*), we focus here only on modeling the binary topology of the connectome. Additionally, while previous work has identified an initial set of connections that act as a seed for the model (*29, 30, 32*), we initiate our models from an empty connection matrix to avoid imposing arbitrary structure on the model network. Another distinction between our implementation and past work is that we used an exponential penalty for the wiring cost term in our models, whereas others have used a power-law form (*5, 30, 32*) or have evaluated both exponential and power-law penalties (*28, 29*). We focus on an exponential penalty for two reasons. First, there is ample empirical evidence that, across different species and resolution scales, the connection probability between pairs of neural elements shows an approximately exponential decay as a function of their distance; the so-called exponential distance rule (EDR) (*19–23, 27, 88*). Our approach thus offers a natural comparison to this extant literature. Second, the scale-invariance of the power-law preserves relative connection distances as a function of global changes in brain size, which precludes an opportunity to study how developmental changes in cortical geometry and associated wiring costs influence connection probabilities in the model, estimated as outlined below.

#### Estimating connection probabilities

The model defined in Eq. 2 is used to determine the probability of forming a connection between two nodes. It is important to note that *θ*_*ij*_ is not itself the actual connection probability, but rather indicates a connection score, such that higher values (indicating more viable connections) are more likely to be formed. The advantage of using *θ*_*ij*_ as a connection score rather than a direct probability is it allows the model to be formulated such that density can be strictly controlled, which is important because many topological properties depend on the number of edges in the network.

Edges are preferentially sampled according to the edge’s own 0 value, divided by the sum of all other possible *θ* values (i.e., the scores for other edges which could possibly be formed), which we term *P*_*ij*_. A single edge is selected at each model iteration according to the probability *P*_*ij*_, and this procedure is repeated until the desired number of edges have been added into the network.

#### Accurately modeling cost-topology trade-offs

The models that we consider here form connections probabilistically and one at a time according to a specific set of wiring rules. The simplest such model that we evaluate considers only spatial factors driven by an EDR,

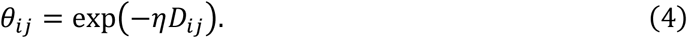

Under this model, connections are formed at random, subject to the constraint that the connection decays exponentially as a function of the distance between two nodes.

Trade-off models commonly studied in the literature include a topological term and have the general form as described in Eq. 1; i.e., *θ*_*ij*_ = exp(−*ηD*_*ij*_) × *T*_*ij*_^*γ*^. The topological term *T*_*ij*_ is intended to counteract the distance penalty imposed by the exponential function if a given connection augments the topological complexity of the connectome. Under this multiplicative formulation of Eq. 1, the influence of the distance and topological terms on *CS*_*ij*_ is modified via a non-linear (power-law or exponential) function. This formulation influences how topology and distance terms interact. In particular, it has the practical effect of ensuring that the topological term only influences the connection probabilities of short-range connections. As an example, Fig. 6A shows the dependence of *CS*_*ij*_ on the connection distance, modelled using an exponential decay for *D*_*ij*_. The parameters for the distance term were selected from results obtained in this paper or previous work (*29*). For a given distance penalty, different values of *T*_*ij*_^*γ*^ only influence the connection score, *CS*_*ij*_, for connections shorter than 55 mm. The effect is exacerbated when using a power-law penalty on the distance term (Fig. 6B). It is thus very difficult for the topological term to overcome the strong penalty on long-distance edges because the topological term is implemented as a non-uniform multiplicative scaling factor across different values of distance. This behavior does not align with a cost-value trade-off, in which the topological value of an edge should counteract its wiring cost, even over long distances. Indeed, interpretation of the model’s parameter estimates is ambiguous due to the complex inter-dependence of model parameters and the fact that they exert two effects in the model: (1) they control the relative relations between different values within a given term; and (2) they control how the different terms scale relative to each other. Because both objectives need to be achieved with the same non-linear function, it is difficult to disentangle the extent to which either is being fulfilled.

**Fig. 6.**
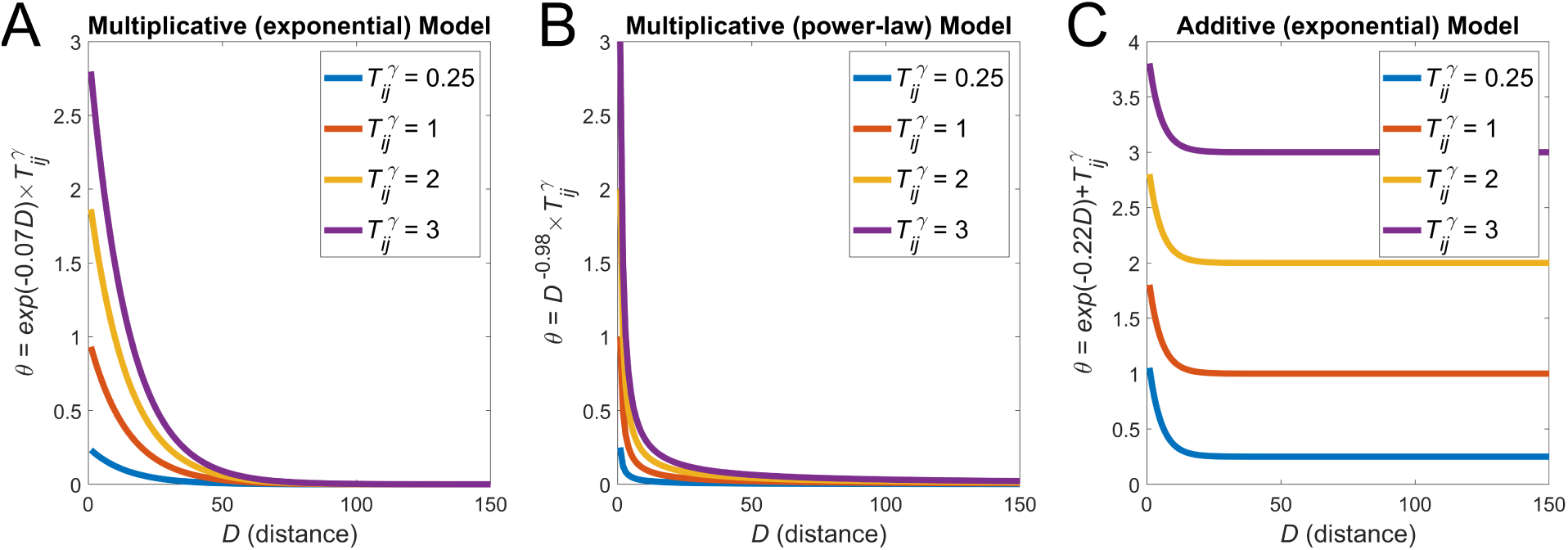
Practical demonstration of multiplicate and additive formulations of the trade-off model. **(A)** *CS*_*ij*_ values calculated over distance with varying values of *T*_*ij*_^*γ*^under a classical multiplicative formulation (e.g., Eq. 1) with an exponential distance penalty. **(B)** *CS*_*ij*_ values calculated over distance with varying values of *T*_*ij*_ ^*γ*^ under a classical multiplicative formulation with a power-law distance penalty. **(C)** *CS*_*ij*_ values calculated over distance with varying values of *T*_*ij*_ ^*γ*^ under our new additive formulation (e.g., Eq. 2) with an exponential distance penalty. For the multiplicative formulation, changes in *T*_*ij*_ ^*γ*^ (which could either be due to changes in *T*_*ij*_ or *γ*) practically only affect short-range connections (**A, B**). Under the additive formulation (**C**), variations in *T*_*ij*_ can influence *CS*_*ij*_ over a broader range of distances.

To avoid these problems, we can formulate an additive wiring rule

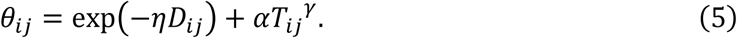

This form allows a single parameter, *α*, to control the importance of *T*_*ij*_ relative to *D*_*ij*_ in determining connection scores. The additive form of Eq. 5 ensures that, for a given value of *γ*, the impact of topology is linear and consistent across all connections, as shown in Fig. 6C over different *T*_*ij*_^*γ*^ and *α* values. Under this formulation, each term can vary independently, meaning that parameters can be selected such that long-range connections can benefit from having a greater *T*_*ij*_^*γ*^ value. This formulation is more consistent with common notions of cost-topology trade-offs, as topology can be sufficiently weighted to overcome the wiring cost of a connection. Moreover, *α* is readily interpretable as the relative weighting assigned to topology vs connection distance in determining connection probabilities, such that higher values indicate a stronger contribution of topology to *θ*_*ij*_.

To interpret *α* as controlling a trade-off between wiring cost and topology in Eq. 5, the distance and topology terms must vary on similar scales. We thus normalize both terms to have a maximum of 1 by dividing each term by its maximum over for all edges that have not yet been added to the network. Our model formulation then becomes

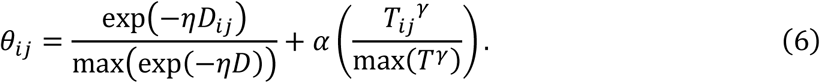

#### Incorporating developmental changes in cortical geometry

Generative models of human brain networks have traditionally only considered wiring costs estimated using physical distances in the adult brain. These ‘static’ models thus neglect the potential impact that developmental changes in brain size and shape, occurring when connections are actually formed, can have on wiring costs. To incorporate these developmental changes, we estimated, for each of the 18 timepoints for which we have fetal scans, a unique inter-regional distance matrix using the ray-tracing procedure described above. When taken with the adult data, this yielded a total of 19 time points. In principle our approach could be extended to include additional time points between birth and adulthood, but we focus here on the prenatal stage because this is when the bulk of inter-regional connections are formed (*77, 78, 89*). We add connections to our model networks in distinct stages, constrained by the corresponding developmental time point, to approximate the effect of changes in brain size and shape. We thus introduce a time-varying wiring cost, *D*_*ij*_(*t*) which indicates the inter-nodal distance between nodes *i* and *j* at timepoint *t*, yielding Eq. (3).

A critical question in this model concerns the rate at which connections should be added to the model at different time points. Detailed empirical data to answer this question are lacking. One study found that expression of GAP-43, a marker of axonal growth, was highly and stably expressed between 21 and 43 weeks post conception, suggesting that this is a period of sustained axonal formation (*89*). Studies of axonal numbers in the developing rhesus monkey have suggested that the number of axons increases linearly during gestation up until birth (*77, 78*). We thus use the simplest possible formulation and add connections at a constant rate at each timepoint *t*, but note that our framework is flexible enough to enable explorations of alternative developmental trajectories.

#### Model evaluation

Model performance was evaluated by comparing the model and empirical node distributions of degree, betweenness, and clustering, and the distribution of connection distances across all edges, as in prior work (Fig. 1D) (*28, 29*). These properties are classical features that are often used to describe brain-network topology. In each case, the distributions are compared using the Kolmogorov-Smirnov (*KS*) statistic, which is quantified as the maximal distance between the empirical distribution functions of two samples, in which lower values indicate greater similarity between distributions (i.e., between the distribution of a topological property in the empirical and model network). Model performance was defined as the maximum *KS* statistic observed across the four-benchmark metrics,

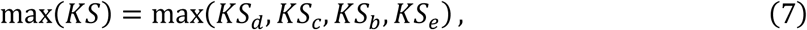

where *KS*_*d*_, *KS*_*c*_, *KS*_*b*_, and *KS*_*e*_ are the *KS* statistic of the degree, clustering, betweenness, and edge length distributions, respectively. In this formulation of model fit, performance is determined by the worst-fitting property. We used the same procedure to assess the performance of each model.

#### Model optimization

To find the optimal values for the parameters *η, γ*, and *α*, in each model for each participant, we used an optimization method developed previously (*29*), implemented as follows:

1. we selected a random sample of 2000 points in the parameter space defined by *η* (evaluated over the range −2 to 10^−10^) and *γ* (varied over the range −8 to 8) and/or *α* (varied between 0 to 8; however when no *γ* was included in the additive formulation, *α* varied over the range 0 to 0.05 for the clu-avg, clu-max, clu-diff, deg-avg, deg-max, deg-diff, and deg-prod models, for all others it varied between 0 to 8). For CGE and MPC models, *γ* was varied over a greater range (CGE: −50 to 250, MPC: 0 to 50).
2. at each point, which represents a specific combination of *η*, and *γ* or *α* values, we generated a network using each of the newly defined parameters (thus making 2000 synthetic networks) and calculate the max(*KS*) fit statistic.
3. once all networks were evaluated, we used a Voronoi tessellation to identify regions (cells) of the parameter space associated with low fit statistics. A further 2000 points in parameter space were preferentially sampled from each cell according to the relative probability 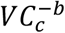, where *VC*_*c*_ is the max(*KS*) of cell *c*, and *b* controls the likelihood with which cells with a low max(*KS*) will be sampled (i.e., a larger value of *b* indicates a greater likelihood of sampling from low max(*KS*) cells).

Steps 2 and 3 were repeated five times, resulting in a total of 10,000 points being evaluated. At each repetition, the probability of sampling cells with better fits is increased (going from *b* = {0, 0.5, 1.0, 1.5, 2.0}), thus converging to an approximate optimum. This optimization was conducted for each model fitted to each individual participants’ network. An advantage of this optimization approach is that it allows for adequate sampling across the entire parameter space to visualize how changes in parameters affect the model and for the identification of a global (approximate) optimum (Fig. 1E).

#### Cross-validation

The one-parameter spatial model has lower complexity than models that include topology, which have two free parameters. Our additive formulation has three free parameters in total. To enable fair comparison across models with varying complexity and to minimize over-fitting, we developed a leave-one-out cross-validation procedure to assess out-of-sample model performance and generalizability. For each participant *s* in our sample of *N* individuals, we generated synthetic networks using the best-fitting parameters obtained for the other *N* − 1 participants. We cross-validated results with respect to the optimal parameters for the other *N* − 1 participants to account for variability across connectomes and to assess out-of-sample performance. For each such parameter combination drawn from the other participants, we iterated the model 20 times to account for variability in the stochastic models. We took the mean fit (of the test statistic, max(*KS*)) across these 20 runs, and then took the average of these means over the *N* − 1 parameter combinations as our cross-validated fit statistic, *F*_*CV*_, for each participant. This approach allowed us to obtain a distribution of *F*_*CV*_ values over participants for each model (Fig. 1E). Unless stated otherwise, all results are reported using this cross-validated fit statistic. While alternative cross-validation procedures are possible, we deemed this leave-one-out procedure to be the most computationally expedient, given the large number of model iterations that were required. To compare *F*_*CV*_ for given models, we used Bonferroni corrected (corrected for 325 tests for comparisons between topological static and growth models; 120 tests for comparisons between physiological models), Wilcoxon signed-rank tests. This non-parametric test was used as *F*_*CV*_ was not always normally distributed.

### 4.7 Modeling brain network topography

According to the procedures outlined above, model fits were optimized for reproducing the statistical properties (node- and edge-level distributions) of network topology. As previously stated, the same distribution may be realized with different spatial embeddings, and it is the spatial embedding or topography that defines the roles ascribed to brain regions, such as which areas are network hubs. We thus sought to quantify the degree to which the models were also able to capture the spatial embedding of the same topological properties used in the model-fitting procedure; i.e., degree, clustering, betweenness, and mean nodal edge length. To this end, we evaluated the Spearman correlation between the nodal values estimated for each property in the empirical and synthetic networks. A high correlation implies that the generative model can accurately capture the relative nodal rankings, and thus spatial embedding, of that particular topological measure.

## Acknowledgments

We would like to thank Richard Betzel for sharing the optimization code for the generative modeling. This work was supported by the MASSIVE HPC facility (www.massive.org.au). Data were provided [in part] by the Human Connectome Project, WU-Minn Consortium (Principal Investigators: David Van Essen and Kamil Ugurbil; 1U54MH091657) funded by the 16 NIH Institutes and Centers that support the NIH Blueprint for Neuroscience Research; and by the McDonnell Center for Systems Neuroscience at Washington University.

## Author contributions

S.O., B.F., K.A., and A.F. all contributed to the experimental design. S.O., A.A., R.S., and C.P., prepared data for the analysis. S.O. conducted the main analysis. S.O. and A.F. wrote the original draft, and B.F., K.A., A., R.S., and C.P., provided feedback.

## Data and materials availability

Data reported in this paper can be downloaded from https://figshare.com/s/9cdeb8854ed51e0eae99 and code to reproduce all analysis reported in this paper can be found at https://github.com/StuartJO/GenerativeNetworkModel

## Supplementary Materials for

### Supplementary Text

To validate our new additive formulation of the cost-topology trade-off (Eq. 2) model, we compared its performance to the classical multiplicative form (Eq. 1). We first considered the additive formulation without the *γ* parameter (i.e., without non-linear scaling of the topology term). Under this additive formulation, 10 of the 12 trade-off models, all of which are based on degree and clustering, show comparable performance to the spatial model. In these 10 trade-off models, *α* ≈ 0, indicating that variations in topology have minimal influence on model performance and that wiring is largely determined by spatial constraints. The two additive trade-off models that perform better than the spatial model correspond to the homophilic matching and neighbor models. Critically, the mean *F*_*CV*_ of the additive matching index models (0.23 ± 0.01) is significantly lower than the multiplicative matching index model (0.27 ± 0.01, *p* < 0.0001; it is also significantly lower than the multiplicative clu-avg model which was the best-fitting multiplicative model on average: 0.26 ± 0.02,), *p* < 0.0001 (fig. S1A). Thus, when considering the best-fitting models, the additive formulation offers a more accurate representation of the data than the multiplicative formulation. Examining the types of edges formed by the empirical and best fitting networks, the additive matching model shows the most notable improvement in capturing mid-to long-range connections when compared to its multiplicative counterpart (fig. S2). This result confirms our intuition that the additive formulation is more effective in trading-off topological value with distance in determining connection probabilities.

We next evaluated whether an additional non-linear scaling of the topology term *T*_*ij*_ improves model performance. To this end, we compared the performance of additive models with and without the scaling exponent *γ*. We found that the non-linear scaling improves the fit of all trade-off models relative to strictly linear variants of the additive models (fig. S1B; the best non-linear additive models are also superior to all multiplicative models, fig. S1C), but the improvement for the best-fitting matching index model was small (*F*_*CV*_ = 0.22 ± 0.01 for the non-linear additive model and *F*_*CV*_ = 0.23 ± 0.01 for the additive model; fig. S1B). The superior performance of the non-linear additive model further suggests that the influence of topology on connection probabilities requires some non-linear scaling, such that regions with a high topology score (or low if *γ* is negative) are disproportionately more likely to form a connection.

**Fig. S1.**
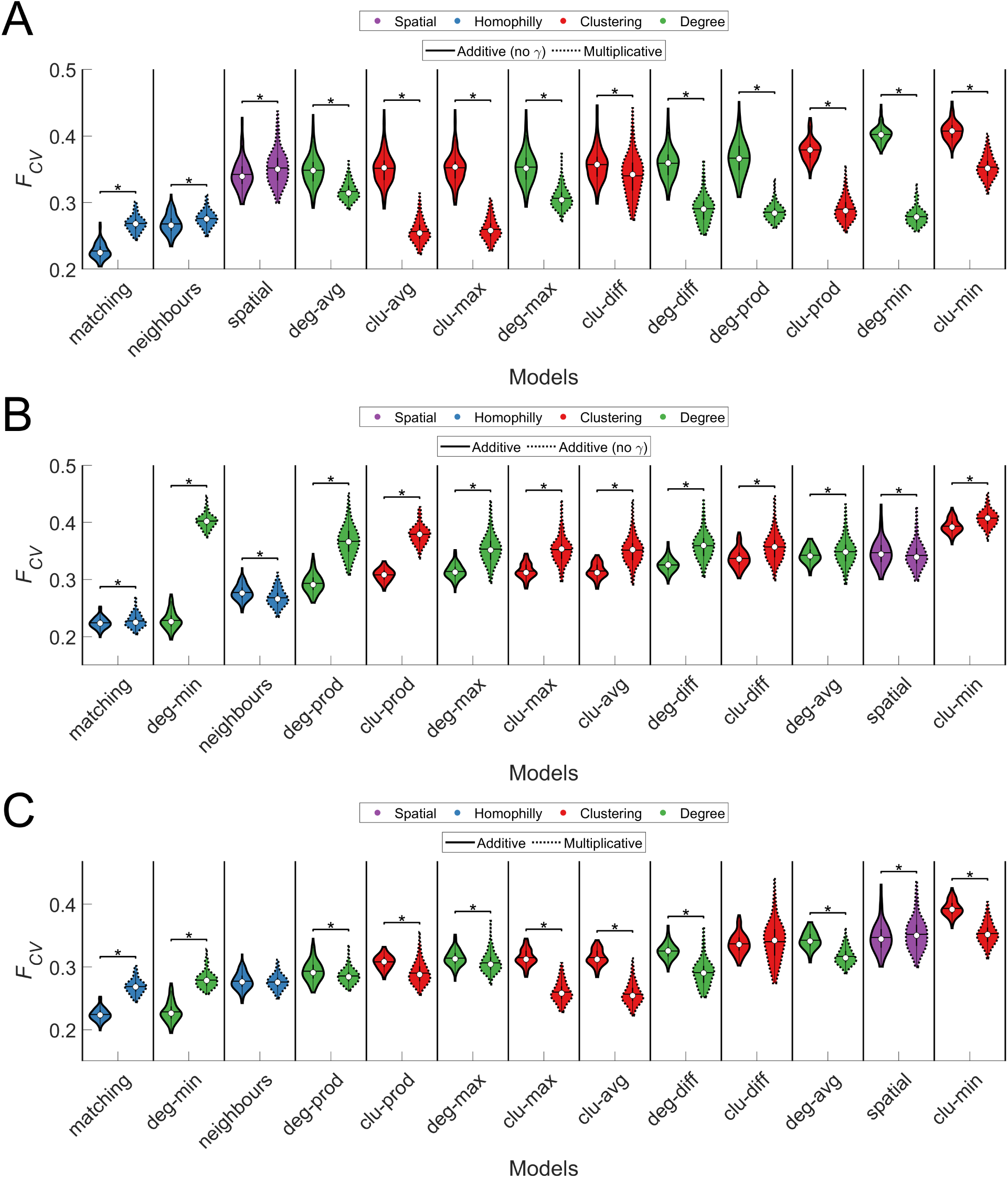
Cross-validated *F*_*CV*_ values for static generative models. The color of each violin plot indicates the topology metric used in the model: homophily is shown in blue, clustering in red, degree in green, communicability in orange, and geometric in purple. The white circle indicates the median of each distribution, while the horizontal black line indicates the mean. **(A)** *F*_*CV*_ values for the additive (with no *γ* parameter) and multiplicative model formulations. Each point in the distribution corresponds to the *F*_*CV*_statistic obtained for one of the 100 individuals in our sample. **(B)** Comparison of *F*_*CV*_ values for the additive and additive (with no *γ* parameter) formulations. **(C)** *F*_*CV*_ values for the additive and multiplicative model formulations. * *p* < 0.05 Bonferroni corrected (325 tests), Wilcoxon signed-rank test. The additive matching model achieves the best performance.

**Fig. S2.**
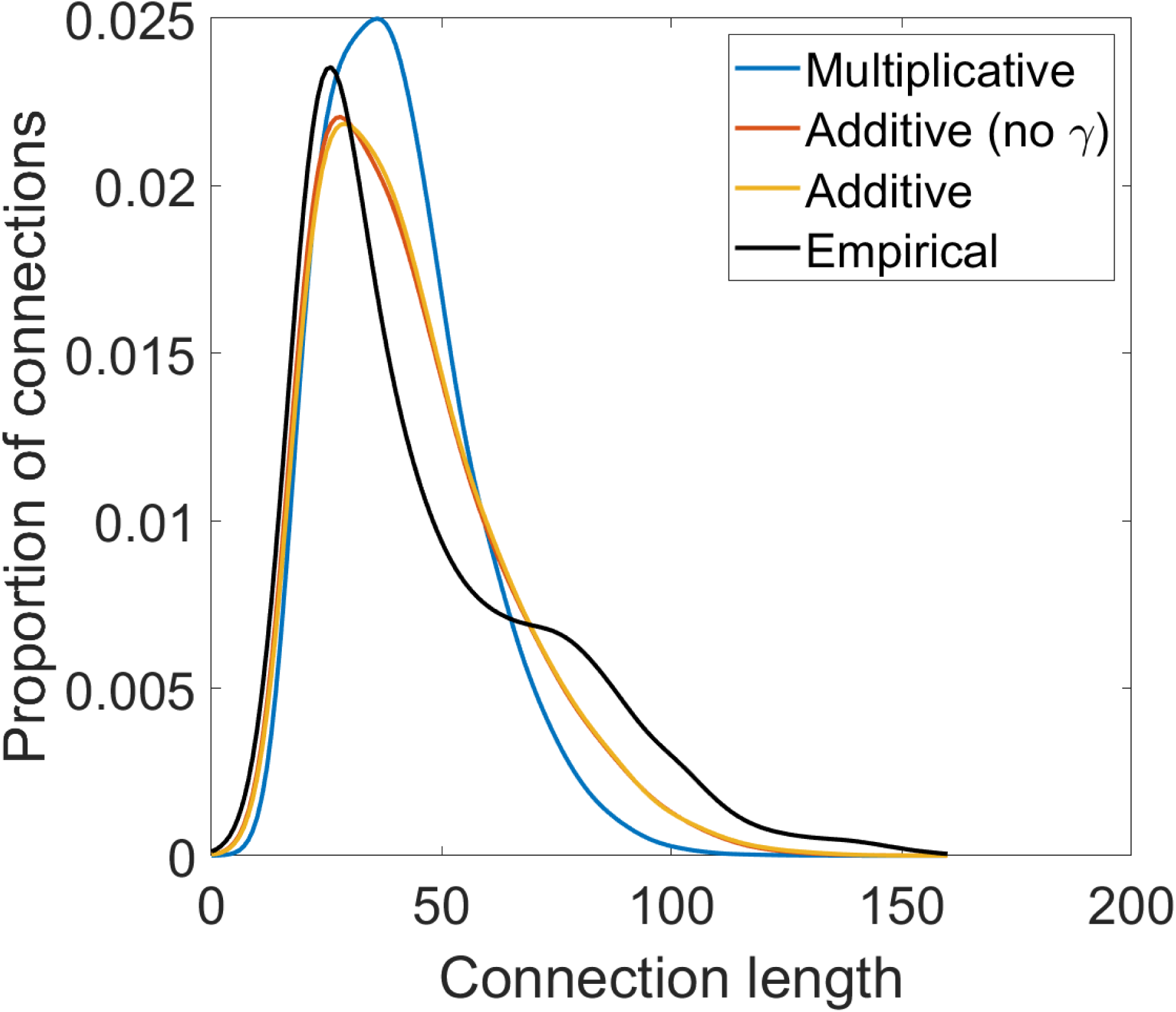
Kernel density plot of connection lengths for matching index models. For the best matching model network for each participant, the kernel density of its edge lengths was calculated and then averaged. This was done for each model formulation. Additive formulations more closely match the empirical data, and are better able to reproduce long-range connections.

**Fig. S3.**
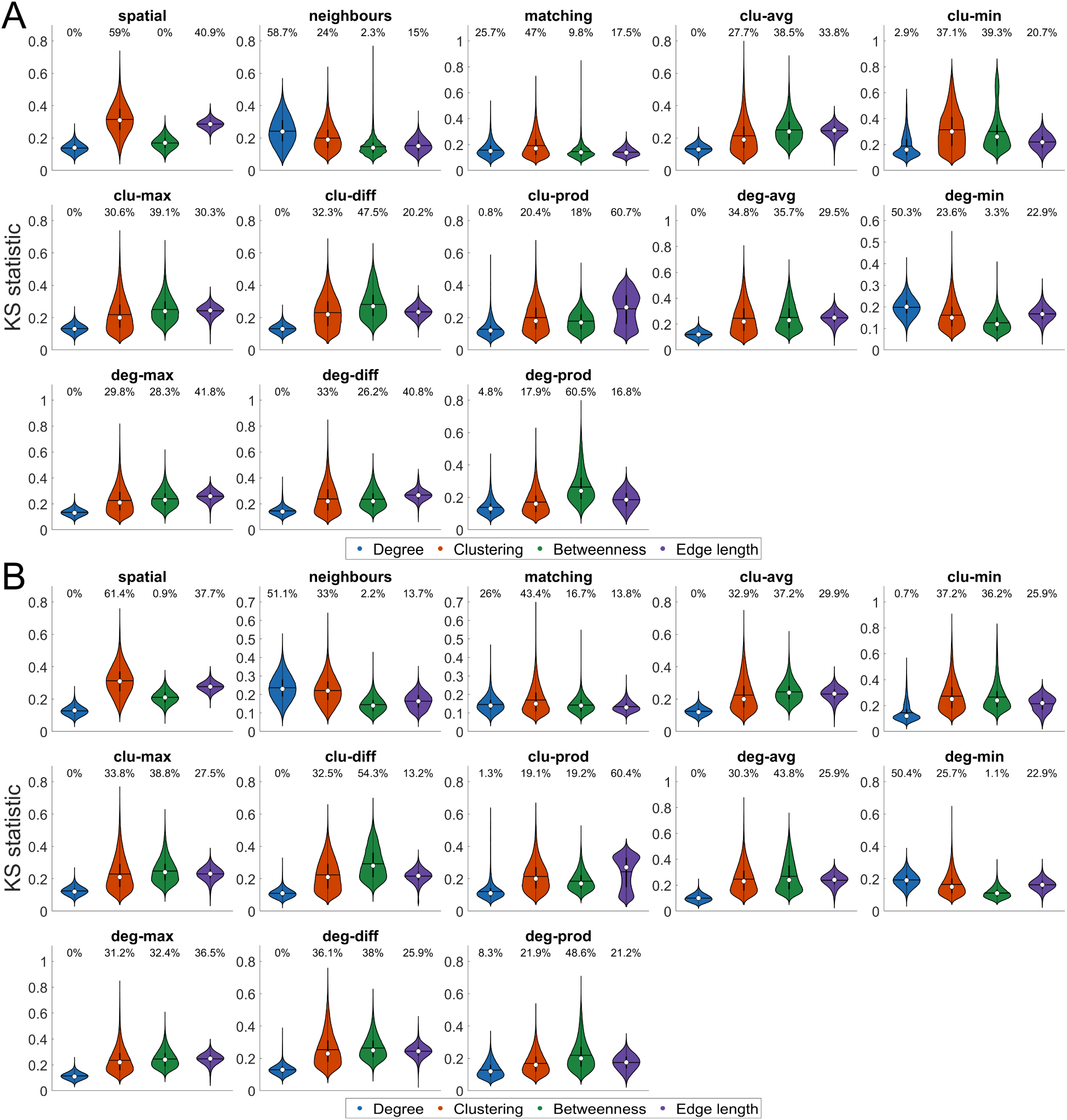
Kolmogorov-Smirnov (KS) statistics for each cross-validated static and growth model network. Each violin plot shows the distribution of KS statistics across all model networks used to derive the cross validated fit *F*_*CV*_ for static (**A**) and growth (**B**) model variants, for each topological feature used in the model fitting procedure; namely, degree, clustering, betweenness, and edge length distributions. The white circle indicates the median of each distribution, while the horizontal black line indicates the mean. The number above each violin plot indicates the percentage of times that respective KS statistic determined model fitness i.e., max(*KS*).

**Fig. S4.**
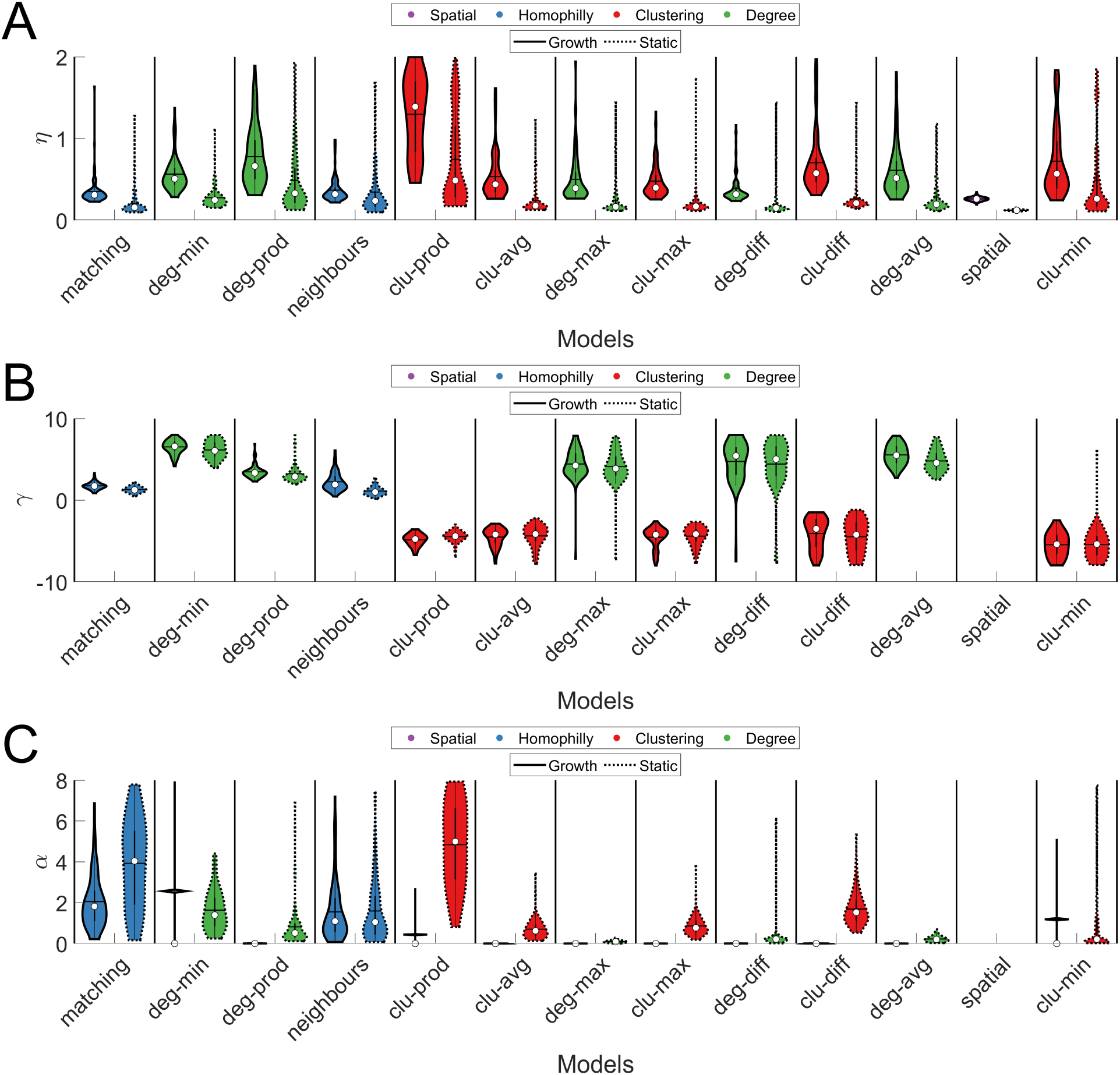
Model parameters of best-fitting static and growth models. **(A)** Values of *η*. Higher values indicate a stronger distance penalty. **(B)** Values of *γ*. Higher values indicate a stronger non-linear scaling of topology, such that high values exert a proportionally greater influence on connection probability. **(C)** Parameters for *α*. Higher values indicate a stronger bias of topology relative to wiring cost on connection probability. The color of each violin plot indicates the class of topology metric used in the model: homophily is shown in blue, clustering in red, degree in green, and spatial in purple. The white circle indicates the median of each distribution, while the horizontal black line indicates the mean. Growth models require a larger *η*, and thus stronger distance penalty, to achieve the best fits.

**Fig. S5.**
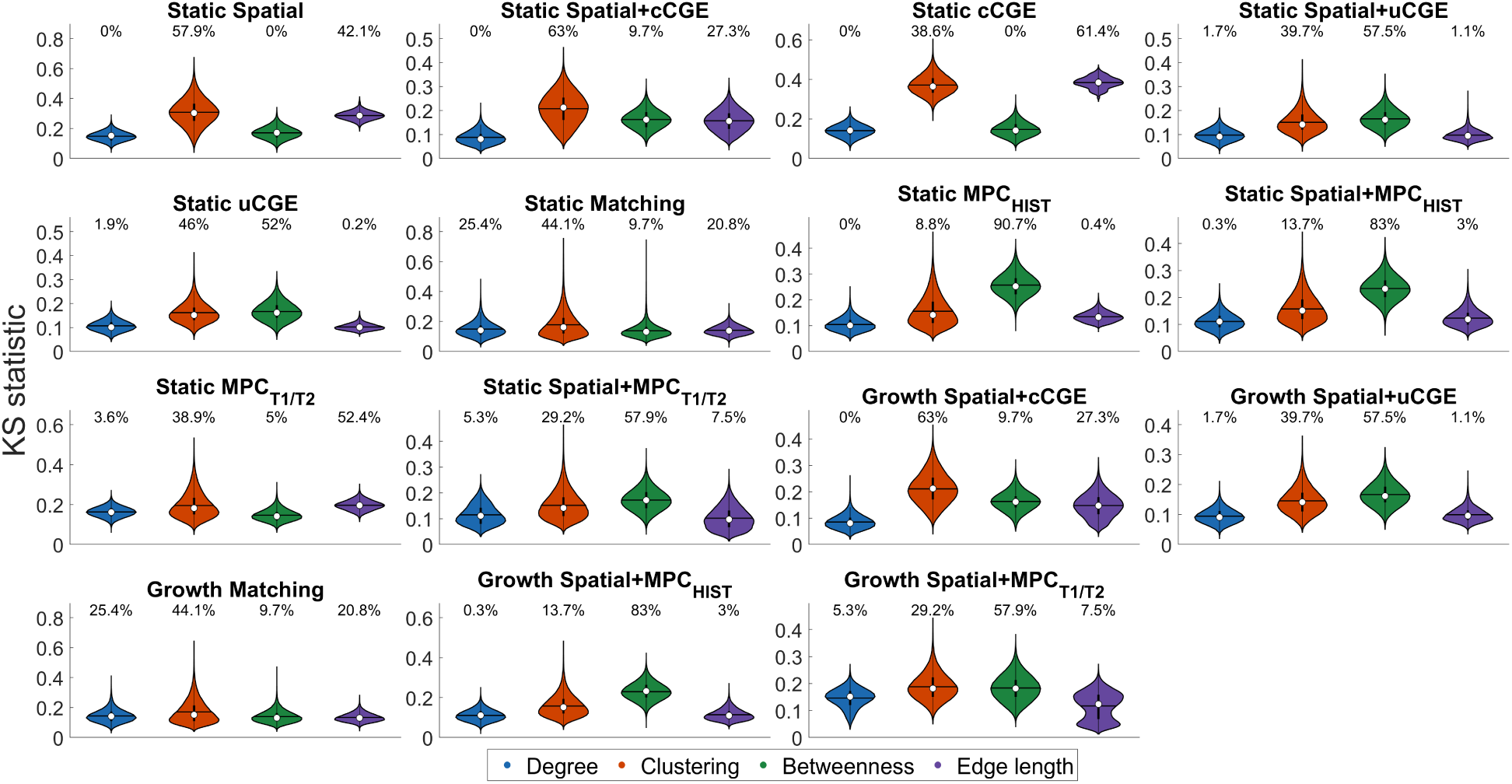
Kolmogorov-Smirnov (KS) statistics for each physiological model network. Each violin plot shows the KS statistic for the relevant topological feature (node degree, node clustering, node betweenness, or edge length) for all networks used to calculate *F*_*CV*_ for each participant. The white circle indicates the median of each distribution, while the horizontal black line indicates the mean. The number above each violin plot indicates the percentage of times that KS statistic determined model fitness i.e., max(*KS*), which itself was used to derive the cross validated fit *F*_*CV*_.

**Fig. S6.**
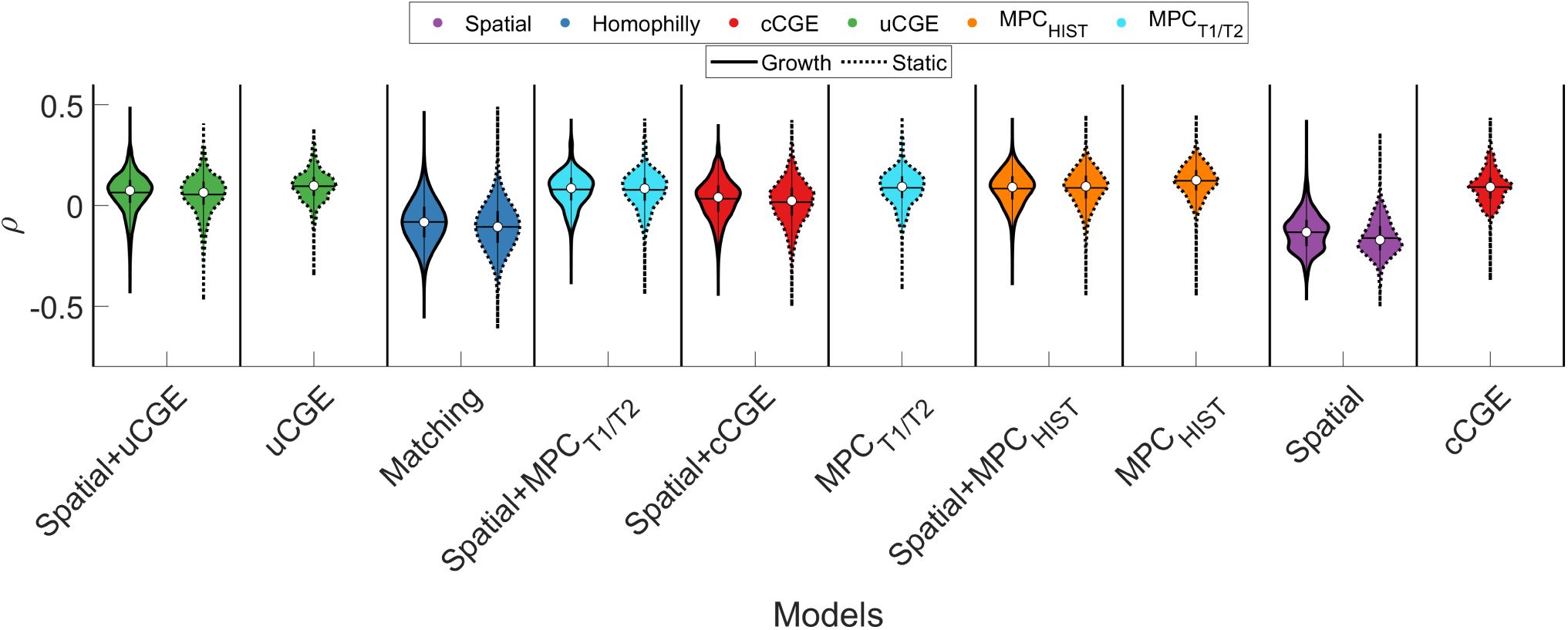
Evaluating model accuracy in capturing degree topography across the full parameter landscape. Each violin plot shows, across all parameters produced during optimization for all participants, the correlation between model and empirical node degree. The color of each violin plot indicates the topology metric used in the model while the white circle indicates the median of each distribution, while the horizontal black line indicates the mean.

**Fig. S7.**
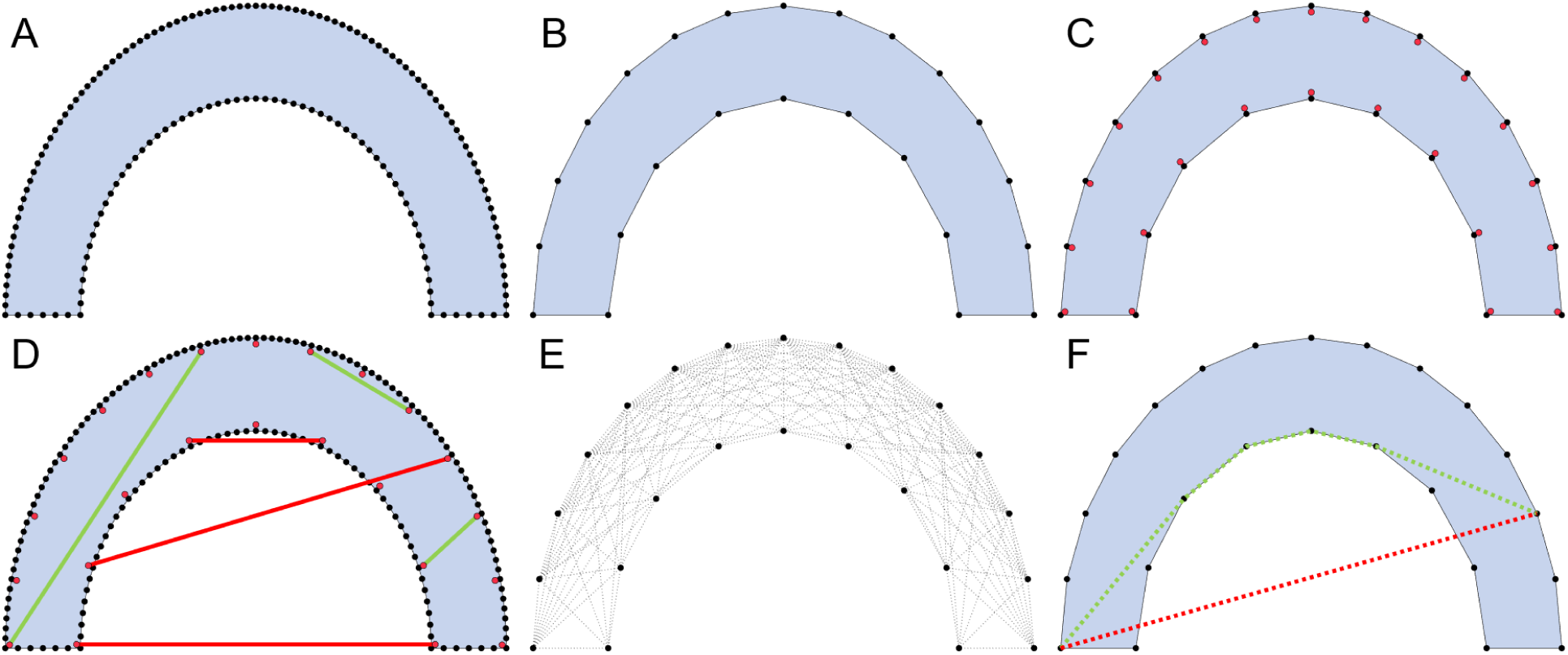
Example of connection distance estimation. (**A**) Simplified example of a cortical surface reconstruction with black points representing surface vertices. (**B**) The surface is downsampled so that only 15% of the original vertices remain. This step minimizes computational burden while preserving the shape of the surface. (**C**) Subsurface points (red dots) are calculated using the vertex normal to avoid precision issues. (**D**) Ray tracing is used to draw line segments between pairs of subsurface points. Green lines show valid trajectories that do not intersect the surface. Red lines show invalid trajectories which are not used for further estimation. (**E**) All pairs of subsurface points are evaluated. If a valid line segment can be drawn between them, then a direct connection (dotted line) is drawn between the corresponding vertices to create a direct connection network. (**F**) Dijkstra’s algorithm is run on the resulting network of direct pairwise direct connections to identify the shortest within-volume distance (i.e., fiber distance) between vertex pairs. Green line shows an example trajectory uses to estimate fiber distance between two vertices. The trajectory used for estimating the Euclidean distance between those same points is shown by the red dotted line.

